# Real-Time Quantification of Cell Mechanics and Functions by Double Resonator Piezoelectric Cytometry — Theory and Study of Cellular Adhesion of HUVECs

**DOI:** 10.1101/2023.01.27.522341

**Authors:** Tiean Zhou, Jingyuan Huang, Lun Xiong, Haibo Shen, Fushen Huang, Wenwei Li, Hange Peng, Zhaohong Su, Weison Pan, Jia Zhao, Zhen Zhou, Dongqin Bao, Linhong Deng

## Abstract

Cell mechanics is closely associated with cellular structure and function. However, the inability to measure both cellular force and viscoelasticity of statistically significant number of cells noninvasively remains a challenge for quantitative characterizations of various cellular functions and practical applications. Here a double resonator piezoelectric cytometry (DRPC), using AT and BT cut quartz crystals of the same frequency and surface morphology is developed to simultaneously quantify the cells-generated forces (ΔS) and viscoelastic moduli (G′, G″) of a population of isolated single cells or cells with different degrees of cell-cell interactions in a non-invasive and real time manner. DRPC captures the dynamic mechanical parameters ΔS and G′, G″ during the adhesions of human umbilical vein endothelial cells (HUVECs) under different ligand densities of adhesion molecules fibronectin or Arg-Gly-Asp (RGD) modified on the gold surfaces of 9 MHz AT and BT cut quartz crystals, and different seeding densities of HUVECs. It is found that both the ligand density and cell seeding density affect the magnitudes of ΔS and G′, G″ and their correlations are revealed for the first time by DRPC. The validity of DRPC is further verified by mechanical changes of the cells in response to treatments with cytoskeleton regulators.

## 1. Introduction

Cell mechanics and mechanobiology are fundamental in development and shape, structure, and function of biosystems, and play important roles in many biological processes. It is well accepted that mechanical force can play as big a role in the eventual shape of a tissue that develops as the genes that are activated during the process. Similarly, cells exert different types of forces to change their shapes and execute complex functions. For some cellular functions, mechanical changes accompany with the biological processes which can be used directly to characterize these functions, e.g., mechanical contraction and relaxation cycles of beating cardiomyocytes,^[1]^ single and collective cell migrations,^[2]^ different phases of cellular adhesion.^[3]^ These physiological functions have important implications in assessing pathological status of major cardiovascular and cancer diseases;^[4,5]^ and drug development and assessments.^[6]^ Mechanical forces affect other cellular functions including proliferation and differentiation which may take days to have functional changes, whereas direct force transmission normally occurs on much short scale of milliseconds to seconds, followed by biochemical signaling in minutes, and cellular responses such as gene expression in hours.^[7,8]^ And the resultant mechanical changes represent a product of various biochemical processes such as cell signaling and gene expression. The other major mechanical parameter of cell stiffness or viscoelasticity is most informative of changes in cellular structure and is tightly associated with cell state and function.^[9,10]^ Moreover, other factors contributing to cellular functions such as soluble hormones and insoluble extracellular matrix would also affect cellular forces and viscoelasticity. Therefore, mechanical changes are as fundamental as genetic and proteomic changes for living cells;^[11]^ however, mechanical information received much less attention as evidenced by the fact that not even a single routine instrument of cell mechanics is available in biological or clinical labs.^[12]^

Mechanical measurements include both active force and passive viscoelasticity-two sides of a coin.^[11]^ Cellular forces are usually measured by measuring the deformations of known elasticities of deformable gels or microposts, the most common one is traction force microscopy (TFM) which belongs to the former. Cell stiffness or viscoelasticity is usually obtained by applying known forces of certain frequencies to living cells and again by monitoring the resulted deformations, atomic force microscopy (AFM) is the gold standard. However, there are some limitations for the current methods. First, most of these methods rely on expensive microscopies to tract deformations which are hard for non-experts to use well. Second, these methods are usually only suitable for single cell tests which are slow and require many tests to obtain statistically meaningful results, which preclude their applications in situations where large amounts of samples are required, such as drug screens. Moreover, cells are naturally in multi-cellular environment, isolated single cells do not mimic the cells’ physiological status, and population of cells of different cell-cell contacts are routinely used in cell biology labs. Third, external forces are usually required to apply to cells to obtain passive viscoelastic parameters which may inadvertently affect cells’ behavior (natural status) and are not suitable for quantitative and continuous characterizations of cellular functions. Fourth, there have been no single method allowing to measure both mechanical parameters of cellular forces and viscoelasticity at the same time impeding the through and complete characterizations of cellular functions, until recently nanoscale elastic modulus and prestress of a single cell were obtained by AFM indentation to generate a cellular mechanome.^[13]^ An ideal system for measuring mechanical changes of living cells should have the following features: 1) inexpensive, easy to operate and measure quantitatively mechanical parameters; 2) able to measure large amount of cells at one time, prefer to have the configurations of conventional cell culture plates to measure populations of cells of different cell-cell contacts; 3) non-destructive, which allows to track dynamic processes from the initial and fast force transmission to long term mechanical changes of various cellular functions; 4) able to simultaneously measure both cellular forces and viscoelasticity, preferably at the most sensitive interface where cells generate active forces. Here we report a new method we term piezoelectric double resonator cytometry (PDRC) which meets most of the above criteria.

The AT and BT cut piezoelectric double resonator technique was previously used for simultaneous measurement of surface stress and mass of rigid materials. ^[14]^ However, it is not clear if it is also applicable to viscoelastic materials such as living cells, as these materials usually change their viscoelastic properties accompanying with the generation of surface stress. AT cut quartz crystal microbalance (QCM) has been widely used to study cell adhesion and viscoelasticity based on simultaneously recording changes in frequency (Δf) and energy dissipation (ΔD) or motional resistance (ΔR) of its oscillating quartz crystals.^[15]^ Surface stress induced frequency change during cell adhesion was previously proposed and measured with AT cut quartz crystals assuming the attached cells as Newtonian fluid drops while the mass effect was neglected.^[16]^ However, such model is mainly applicable to suspended cells;^[17]^ moreover, the single AT cut crystal technique is unable to determine the direction of the surface stress, and BT cut crystals have not been applied for the study of living cells at all.

### 2. Theoretical background and equations for DRPC

The silicon dioxide quartz crystals are anisotropic in that their physical properties are different along different molecular axes of the crystal. The major axis of quartz growth is called the optic axis and is labeled the Z axis in an orthogonal X, Y, Z coordinate system where the Y axis is the mechanical axis and the X axis is the electrical axis (Figure 1A).

**Figure 1.**
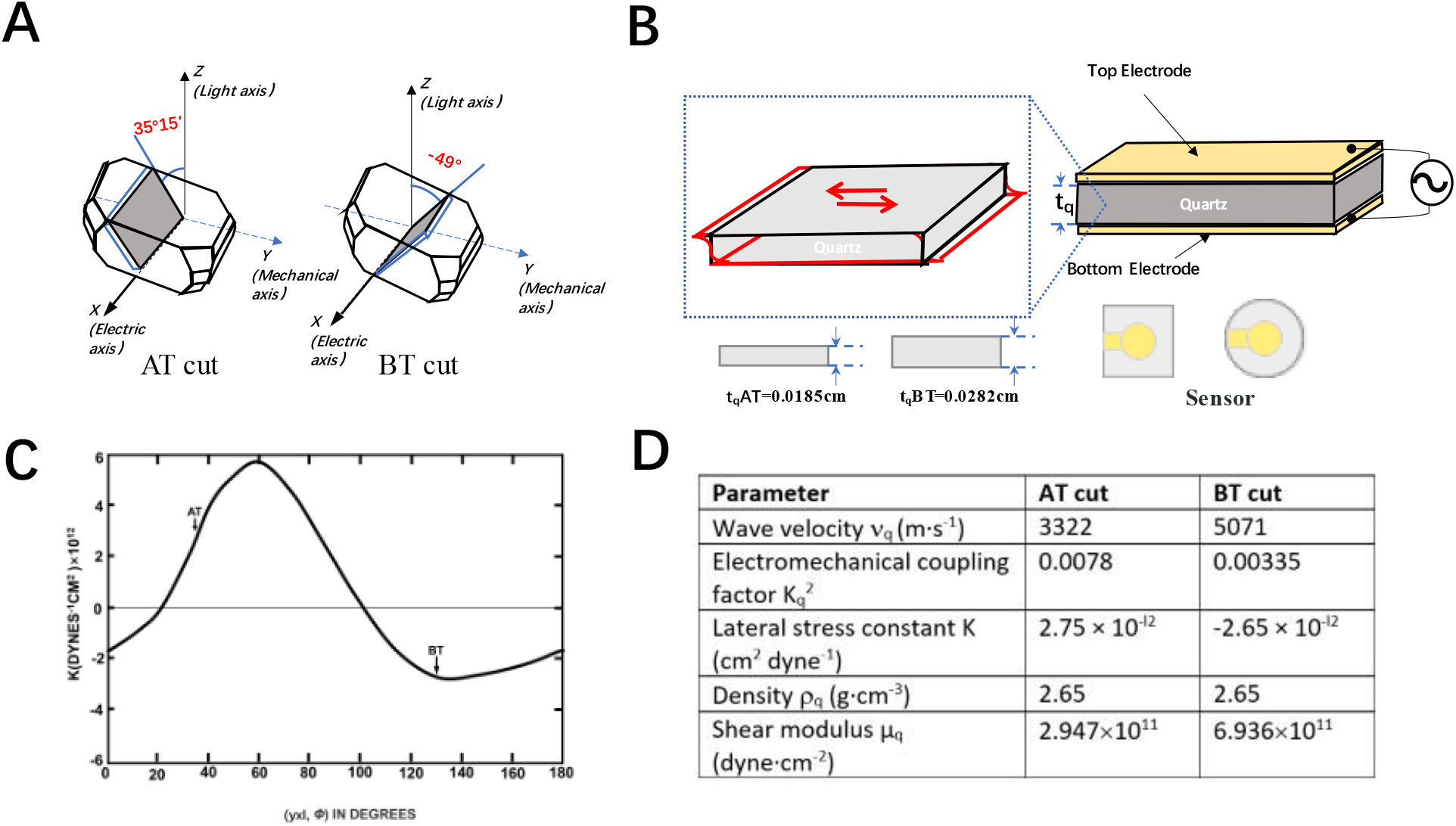
Schematic illustrations of AT and BT cut crystals (A), their thickness shear mode (TSM) oscillation under an AC electric field (B), lateral stress constant as a function and Y-rotated cut angle (C), physical parameters of AT and BT cut crystals which determine their mechanical sensitivities to mass, viscosity, viscoelasticity and lateral stress exerted on the crystals.

A slice of blank quartz used for making a resonator is obtained by cutting the crystal at specific angles to each axis. The choice of axis and angles determine the physical and electrical parameters for the resonator. The widely used quartz crystal microbalance (QCM) is the high frequency thickness shear mode (TSM) resonator which can be excited to oscillate at its natural and fundamental frequency at an AC electric field (Figure 1B). TSM is secured by the rotated Y-cut family of cuts corresponds to a rotation of the y-axis about the crystallographic x-axis. In standard notation, the orientation of a singly rotated Y-cut is typically described by (yxl)Φ. The first and second letters describe the directions of the two important dimensions of thickness, length relative to the capital axes, before rotation. The third notation specifies the direction of rotation relative to the plate dimensions length (l), width (w) and thickness (t) respectively. he angle of rotation Φ is positive for counterclockwise rotation. A light change in the orientation of a quartz crystal plate relative to the crystallographic axes does not alter the mode of resonance. However, the effects of temperature and stress to the resonant frequencies are found to be highly sensitive to the crystallographic orientation.^[14,18]^ The two most important cuts, and cuts, their rotation angles are 35°15’ and -49°respectively (Figure 1A), providing a zero-temperature coefficient of frequency at room temperature of 25 °C.^[18]^ However, their lateral stress constants K^AT^ and K^BT^ are almost in the same magnitude but opposite in sign (Figure 1C), this constant and other physical parameters of AT and BT cut crystals are summarized in Figure 1D. The smaller frequency constant of AT cut compared to BT cut makes BT cut crystal thicker for the same frequency crystal and their thicknesses at 9 MHz frequency is given in Figure 1B.

The initial and most popular application of QCM is for small mass sensing based on the so-called Sauerbrey equation (1) describing the linear relation between changes in the resonator mass and in the resonance frequency. Where Δm is areal density as the mass per unit area. Equation (1) is valid for thin, rigid and uniform films; however, it is also applicable for non-homogeneous or discontinuous films and the QCM measure the sample’s area-averaged mass per unit area.^[19]^

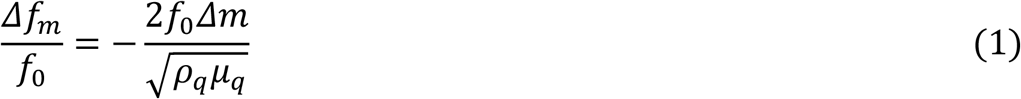

In some cases, surface stress builds in the thin film on the quartz resonator surface which would produce a mechanical bias to the quartz crystal leading to frequency changes of the resonator.^[20, 21]^ EerNisse derived the equation (2) which relates the fractional change and the integral through the film thickness of the lateral stress S_f_, the force per unit width acting on the quartz.

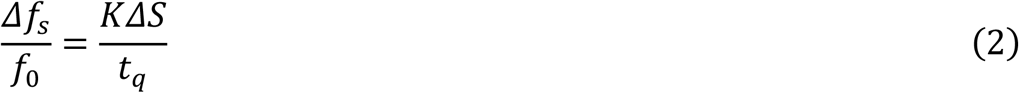

Where K is the lateral stress constant for a given crystallographic orientation of the quartz crystal (Figure 1C).

Equation (2) was derived for stresses built in a thin film on a quartz resonator with the assumption that the lateral stress is isotropic based on the theory of Thurston and Brugger for small-amplitude acoustic wave propagation in a homogeneously stress-biased medium.^[20]^ Quartz itself is anisotropic so there are some differences between the results of the isotropic formulation of equation (2) and the literature value of K, however, the differences are generally less than ten percent.^[22]^ Moreover, cell adhesion induced surface stress was measured by AT-cut quartz crystals and it was demonstrated through the combination of scanning electrochemical spectroscopy (SECM) that the average force induced frequency/motional resistance ratio (stress constant) is about 120±7 Hz·Ω^-1^over different locations of the sensing electrode indicating its approximately homogeneously feature.^[16]^

As far as the film side, though it was assumed that the film should be also isotropic and homogeneous, but in reality, the samples could be heterogeneous both perpendicular and parallel to the QCM surface as identified that the initial nucleation process during film deposition could also induce intrinsic stress and the stress level is typically highest for the first few monolayers of a thin film deposited onto to the substrate, but then normally decreases as subsequent monolayers are deposited;^[22]^ therefore, the average values or ranges of average values of the integral of the lateral stress over the thickness of the thin film are given by equation (2).

When in liquids, liquid’s viscosity and viscoelasticity would induce not only change in crystal’s frequency, but also damping (energy dissipation) as marked by increased motional resistance or bandwidth of the crystal.

The strict relations between the measured changes in frequency (equation 3), motional resistance (equation 4) of AT-cut crystals and the liquid viscosity (η), density (ρ) were derived by Kanazawa and Gordon,^[23]^ Martin et al.^[24,25]^

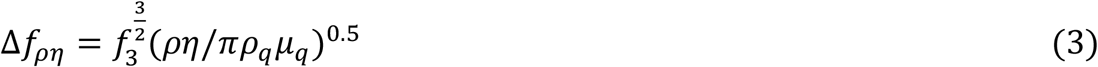

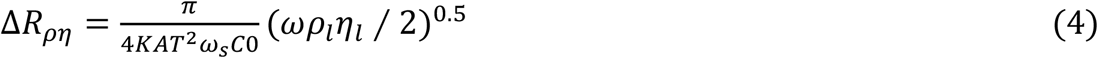

Where C_0_ in equation (4) is the static capacitance of the crystal.

Based on the load impedance and the small-load approximation when Δf/f_0_<<1,^[26]^ a semi-infinite viscoelastic medium induced relative change in complex frequency shift Δf^*^ is given by

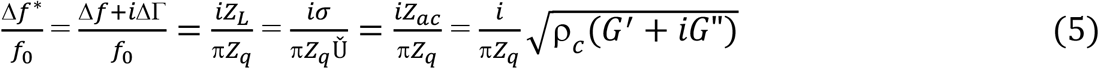

Δf* =Δf+iΔΓ is the complex frequency shift; Δf, ΔΓ are the shifts in frequency and bandwidth. Z_L_ is the load impedance which is defined as the ratio of stress σ and particle velocity Ǔ (speed), and is equal to the acoustic impedance of the medium Z_ac_ if the load is a semi-infinite viscoelastic medium

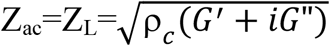

ρ_c_, G′, G″ are the density, storage modulus and loss modulus of the medium. 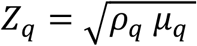 is the acoustic impedance of quartz crystal. For complex material such as living cells, vesicles, the stress-speed ratio can be replaced by an area average stress-speed ratio to calculate the area average load impedance Z_L_ through the measured changes in frequency and bandwidth. The relative frequency shift induced by a semi-infinite viscoelastic medium is

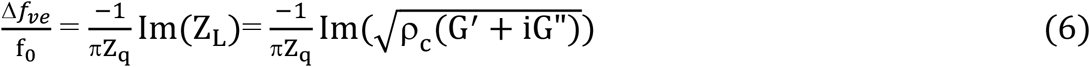

The Sauerbery equation 1 and the above theoretical relations between the measured changes in frequency, motional resistance or bandwidth and the liquid’s viscosity, viscoelasticity were derived for AT-cut quartz crystals, however, it was theoretically predicted that equations (1) and (3) can be extended to other pure shear mode cuts including BT-cut but with some different response slopes owing to their difference in shear modulus,^[18]^ shear wave velocity or acoustic impedance. Here we further assume that equations (4-6) can also be applied to other pure shear mode cuts including BT-cut.

For isolated cells without cell-cell interactions, during the adhesion of a population of cells to the electrodes of the quartz crystals, F-actin structures and myosin motors are key generators of the forces, cells mainly generate tensile stresses toward the centers of the cells, reflecting to the contractile traction force at the electrode/cells interface. Moreover, F-actin polymerization alone can produce pushing forces.^[27]^ The magnitude and direction of the total stresses generated at the cells-electrode interface are determined by the two opposite traction and polymerization forces (Figure 2). The total stress generated by a population of isolated single cells exerting on the AT or BT cut chip is proportional to the number of the cells. Generally speaking, traction force is much larger than the polymerization force, it is this contractile stress that will change the resonant frequency of the quartz resonator. However, before the formation of focal adhesion and establishment of stress fibers, there is a possibility that the polymerization force could be larger than the traction force so that a net tensile stress would occur within the quartz crystal.

**Figure 2.**
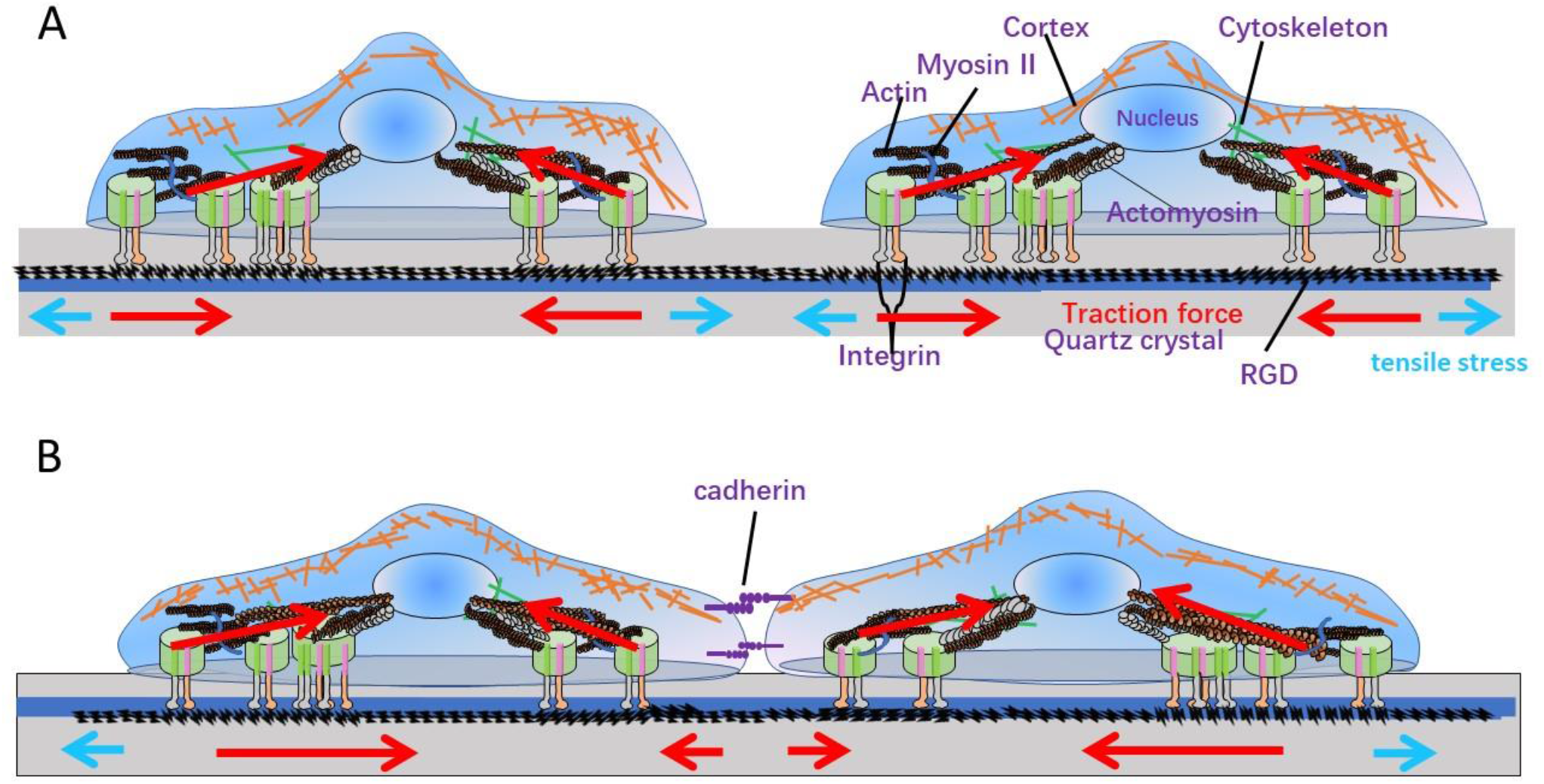
An illustration of the generation and transmission of cell traction force and polymerization force (tensile stress) to the quartz crystal by a population of isolated single cells (A) and cells in contact (B).

When a quartz crystal of fundamental frequency f_0_ is used as the substrate for cell adhesion, the frequency change (Δf) of the crystal relative to its stable value in cell culture media is determined by the following three factors of the living cells: stress exerted (Δf_s_), mass deposited (Δf_m_) and viscoelasticity of the cells (Δf_ve_). To apply and further derive the theoretical equations by using the AT- and BT-cut double resonator quartz crystals to quantify the living cells’ mechanical parameters (generated surface stress and viscoelasticity), the following assumptions are made.

1. As the load impedance is much smaller than the acoustic impedance of the quartz obeying the small-load approximation Δf/f_0_<<1,^[26]^ the frequency changes due to the stress effect, the mass and viscoelastic changes of the cells can be considered as linear perturbations; the total observed frequency change Δf/f_0_ is given by

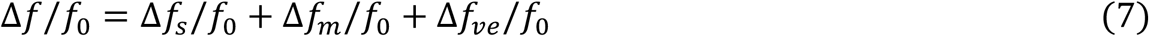

The above equation is justified and addable frequency effects of mass and stress, ^[14]^ mass and viscous loading or mass and viscoelastic loading;^[25,28]^ mass, lateral stress and viscous loading have been demonstrated in previous studies.^[16]^

2. EerNisse equation (2) is applicable for characterizing lateral stresses generated by living cells of confluent, sub-confluent monolayers, cell populations of different cell-cell contacts and isolated cells. Initially, the AT and BT cut double resonator technique was used for measuring the stress effects in thin films that result from film deposition, sputtering, ion implantation in thin films, and the stresses are isotropic and homogeneous only when the film thicknesses reach certain values, the initial nucleation process occurring before the growth of the films on the crystal surface is neither homogeneous nor continuous. Later, it was extended for measuring stresses during lithium transport through the RF sputter-deposited Li_1-δ_CoO_2_ film, ^[29]^ hydrogen adsorption/absorption on Pd overlayers,^[30]^ pH-dependent phase transitions of self-assembled thiol monolayers with carboxylic acid functionality,^[31]^ and even laser illumination resulting in thermally induced radial stress.^[32]^ The stresses exerted on the quartz crystals accompanied with these processes may not be isotropic and homogeneous, in particular, thermal shocks are generally non-homogeneous, however, the double resonator technique is still able to measure the total stress or the areal average stress exerted on the quartz crystals. Vertically, the stress along the thickness direction of the film is also not homogeneous.^[22]^ Therefore, the EerNisse equation can be used to measure not only the lateral stress generated by continuous and homogeneous cell monolayer; but also the total or the areal average lateral stress generated by subconfluent monolayer, cell populations of different cell-cell contacts as well as isolated cell population.

3. Because the decay length of the thickness shear wave of quartz crystals working at MHz range (penetration depth to the living cells) is much smaller than the thickness of the cells, for a dense layer of cells on the sensor surface, the living cells can be considered as a semi-infinite viscoelastic load.^[33]^ For non-continuous, heterogenous cell population of different cell-cell contacts or isolated single cell population, the area average load impedance Z_L_ can be used to describe the cells’ viscoelasticity.^[26]^ Moreover, it is assumed that the cells are homogeneous in viscoelasticity; and the viscoelasticity induced relative frequency change can be described by equation (6).

4. As we always measure the relative changes of cells loaded crystals comparted to the cells-free crystals, the contributions of the surface morphology, surface coating etc. of the crystals can then be subtracted; the measured relative changes in QCM’s frequency and motional resistance are meanly representing the pure contribution of the living cells, so that the cells’ generated forces and viscoelasticity can be precisely quantified by the measured changes in QCM’s parameters.

5. The AT and BT cut crystals have the same morphology (roughness) modified with the same surface material or molecule in the same way, and the same batch and number of cells are added to both AT and BT cut crystals so that they would adhere almost the same living cells and the cells are in almost the same status on both the crystals.

With the above assumptions, apply equations 1, 2, 6 to equation 7 and equation 7 can be re-expressed as:

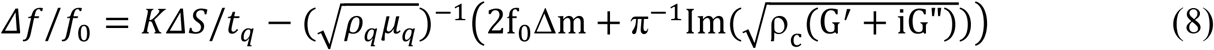

Specifically, for AT cut crystal:

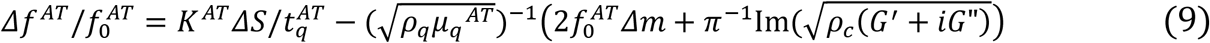

For BT cut crystal:

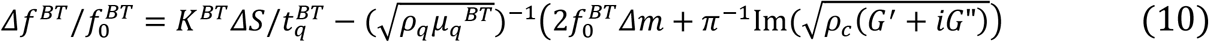

Consider the relationship between the thickness of quartz crystal and frequency, elastic modulus.

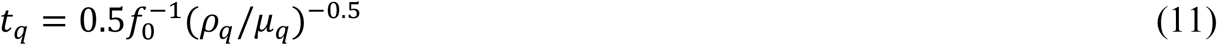

Assume that the frequency of AT cut quartz crystal equals to that of the BT cut crystal, f_0_^AT^=f_0_^BT^=f_0_ and combine equations 9-11, the contributions of the cells’ mass and viscoelastic moduli to the frequency shifts in the parentheses of the second terms in equations (9) and (10) can be cancelled out; hence we obtained equation (12) to quantitatively measure the surface stress ΔS generated and exerted to the quartz crystal by a population of living cells in real time t.

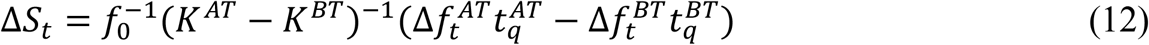

This equation is different from the original EerNisse equation in that both the AT cut and BT crystals must have the same frequencies, which is a special case of the EerNisse equation to make it applicable to the measurement of surface stress generated and exerted by a dynamic viscoelastic load as living cells.

Where K^AT^ =2.75×10^−l2^cm^2^ dyn^-1^, K^BT^ = -2.65×10^−l2^cm^2^ dyn^-1^ are constants; t_q_^AT^ and t_q_^BT^ are the respective quartz crystal thicknesses in cm, f_0_ is the original resonant frequency of AT and BT cut crystals in Hz before cells are added, Δf_t_^AT^,

Δf_t_^BT^ are the frequency shifts given by the new resonant frequencies after the resonators experience the surface stress effects minus the respective f_0_ at time t. The above formula is universal and suitable for the accurate measurement of surface stress or traction force generated by interface material (including cells) accompanied with changes in the material′s mass and viscoelasticity. ΔS>0 indicates that the cell exerts compressive stress (traction force) to the QCM electrode while the cell generating intracellular tensile stress towards its center, and ΔS<0 indicates that the cell exerts tensile (protrusive) forces to the QCM electrode through actin polymerization (Fig. 2A). Thus, the stresses exerted to the quartz substrates would induce measurable changes in QCM frequencies of AT and BT cut crystals, which determine the magnitude and direction of ΔS and the relative contributions of the contractile and protrusive forces exerted by the cell.

When f_0_^AT^=f_0_^BT^=9 MHz, equation 10 can be further simplified to

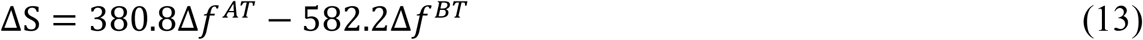

The units for Δf^AT^、Δf^BT^ are Hz, and the unit for ΔS is dyne/cm。

The above eq. 12 was derived assuming no cell-cell interaction and is therefore suitable for quantitative measurement of intracellular stress or protrusive force generated by isolated single cells. However, as the increase of cell number or seeding density, the distance between cells decreases to a certain extent, at the cell-cell contact would produce intercellular contractile stress opposite to the single cell tensile stress, thereby reduce the magnitude of cells’ traction force at the cell/sensor interface (Fig. 2B). As further increase of the cell-cell interactions, cell-substrate contact (spreading) area would decrease and the cells generated stress would change from tensile to contractile, resulting in dominated tensile stress in the quartz crystals changing the sign of ΔS from positive to negative. By calculating the magnitude and direction of the average force exerted by a single cell on the DRPC chip, the degrees of cell-cell interactions and the compactness of the cellular monolayer can be revealed.

The frequency variations caused by the exerted stress ΔS on the DRPC chips can be calculated by the following formula:

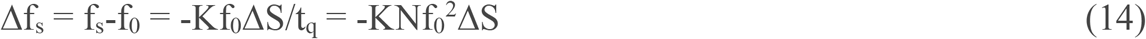

Where fs and f_0_ are resonant frequencies of the AT or BT cut chips subjected to the stress and free of the stress respectively; K, N are stress and frequency constants which are cut dependent. Therefore, stress induced frequency shift is proportional to the square of the resonant frequency of the crystals.

Quantitative measurements of living cells′ viscoelastic parameters using QCM technique are scarce, aging-related viscoelastic variations of tendon stem cells were characterized by QCM through the measured admittance spectrums [34]. Here we consider the living cells as a semi-infinite viscoelastic load because the decay length of the thickness shear wave of quartz crystals working at MHz range is much smaller than the thickness of the cells. Moreover, when the quartz crystals are modified with cell adhesion molecules such as RGD or fibronectin, the cells-sensor distance can thus be neglected. Hence, the storage modulus G’ and loss modulus G ″ of a homogeneous monolayer or a dense layer of living cells can be extracted through the measured changes in frequency Δf and half bandwidth ΔΓ based on equation (5) by comparing the real and imaginary parts of both sides of this equation respectively. ^[26]^ ΔΓ = ΔR/4πL_q_, and L_q_ is the motional inductance of the resonator in cell culture media, hence we have:

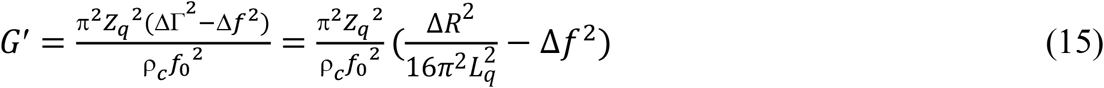

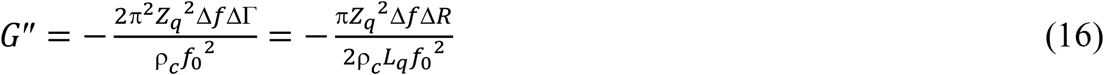

ρ_c_ is the density of the cells which is assumed to be the same of pure water 1.00 g/cm^3^,^[34]^ Δf and ΔR are changes in frequency and motional resistance induced by a homogeneous viscoelastic layer relative to air. In reality, what we measure are the cell adhesion induced changes in frequency and motional resistance relative to cell culture media.

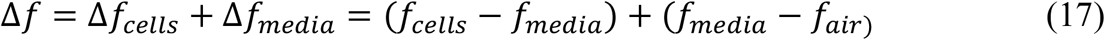

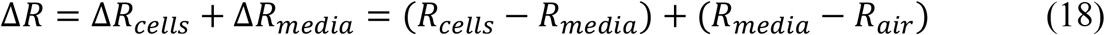

After neglecting the weak mass-effect contribution,^[16]^ the cells’ viscoelasticity induced frequency changes are obtained by subtracting the cells’ stress induced frequency shifts from the measured frequency changes. The frequency and motional resistance of quartz resonators in media can be considered as the same as in pure water at 37°C. Their changes relative to air can be theoretically calculated based on equations (3) and (4).

The applicability of equations (3, 4) and (15, 16) to both AT and BT cut crystals was experimentally verified in both Newtonian sucrose solutions and non-Newtonian (viscoelastic) PEG 2000 solutions at different concentrations (Figure S1).

## 3. Results and Discussion

### 3.1 Effect of surface ligand density on cell adhesion and cytomechanical dynamics during adhesions of HUVECs

Figure 3 shows the time courses of QCM responses during adhesions of 20,000 HUVECs to QCM surfaces coated with FN created at the concentrations ranging from 0 µg/mL to 40 µg/mL and, corresponding cells generated stresses, storage and loss moduli of the cells. It can be seen that the cells induced the largest frequency shifts and motional resistance changes when the surface was coated with FN at intermediate concentration (20 µg/mL), indicating strongest cell adhesion to the surface. Accordingly, the cells generated the largest traction force accompanied with the largest storage and loss moduli, obeying the cellular tensegrity model.^[35]^ Similar trends were observed in HUVECs during adhesions to QCM surfaces coated with RGD created at the concentrations ranging from 0 µg/mL to 100 µg/mL, with largest QCM responses and mechanical parameters occurring when the surface was coated with RGD at 50 µg/ mL (Figure S2).

**Figure 3.**
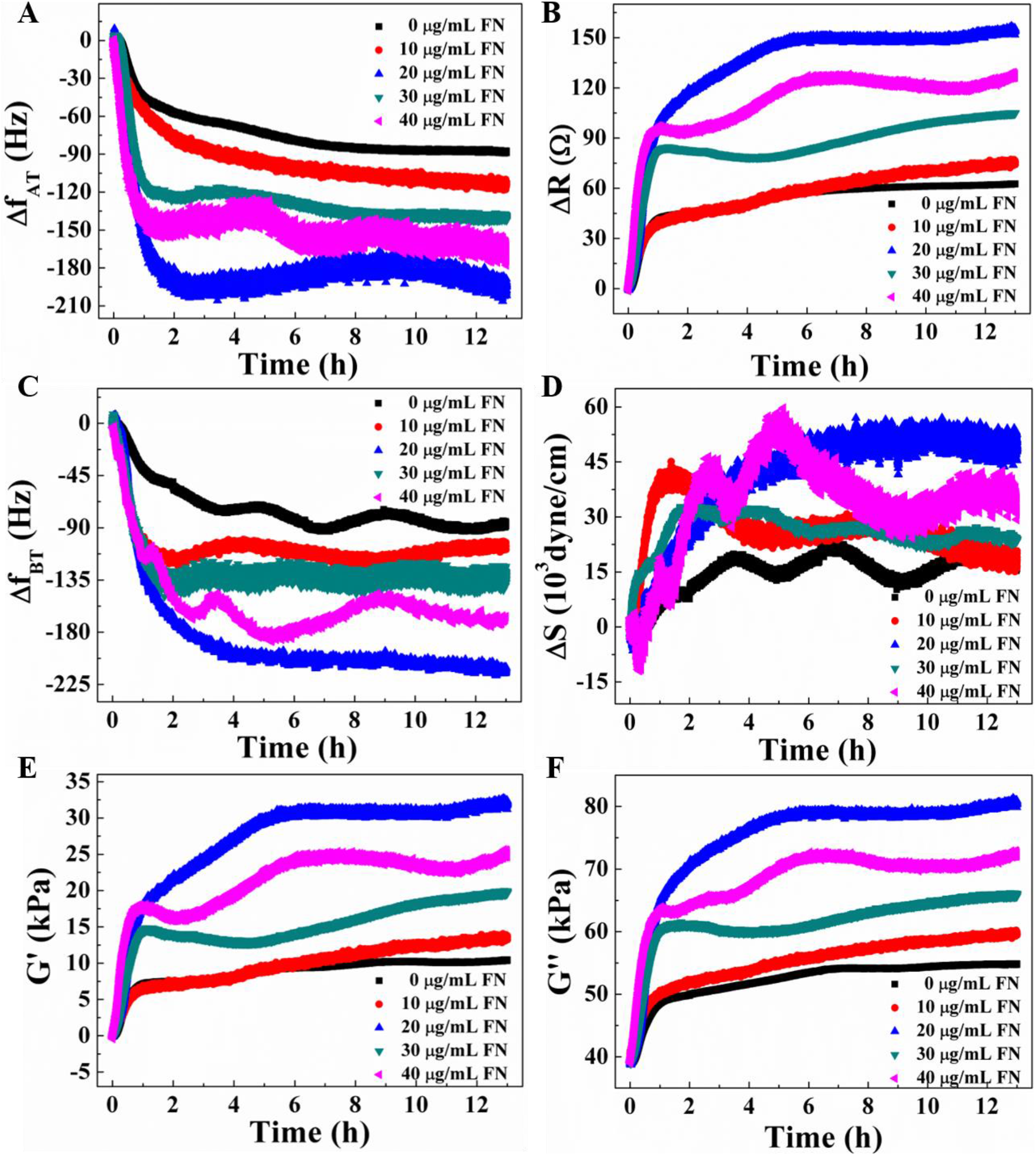
Dynamic QCM responses and cytomechanical parameters captured during the adhesions of 20,000 HUVECs on FN modified Au surfaces created from different FN concentrations. (A, B): AT cut QCM frequency shifts and motional resistance changes; (C): BT cut QCM frequency shifts; (D): Changes in cells′ generated stresses; (E, F): Changes in cells′ storage modulus and loss modulus.

The dynamic behaviors of HUVECs during adhesion can be generally divided into a brief early period of relatively slight changes followed by a rapid and gradual increase, then a final and relatively stable period of the cellular mechanical parameters. In these three phases, the cell generates either contractile traction force that should be directly proportional to the cells’ storage and loss moduli according to the cellular tensegrity model if the actomyosin contractile force dominates, or protrusive force that should also increase with the storage and loss moduli if the force due to actin polymerization dominates. Since both the contractile and protrusive forces lead to cell spreading, it is expected that cell spreading should always be associated with increase of viscoelastic parameters in the cell. In earlier QCM-based study of living cells, it was found that the frequency shift due to cell adhesion is in direct proportion to the cell′s surface coverage or spreading area.^[36]^ Therefore, here we use frequency shift subtracted from the stress induced part (Δf_A_) to indicate the dynamic spreading area of the cells. We indeed found that the viscoelastic moduli (G′, G″) are always in directly proportional to Δf_A_ when QCM surfaces were modified with either FN or RGD created from different FN or RGD concentrations (Figure S3, Table S1). To better understand and distinguish different phases in HUVECs spreading, we did double logarithmic plots of all the parameters including Δf_A_, G′, G″, ΔS over time (Figure 5, Table S2).

Now we can clearly distinguish the following three different phases as identified in: ^[37]^ 1) a brief lag time phase, 2) a fast-continuous spreading phase and 3) a slow spreading phase. The lag time decreased with the increase of FN concentration, being consistent with the previously reported spreading initiation results by FN,^[38]^ whereas no significant difference in lag time was found when RGD concentration is equal to or larger than 50 µg/mL. During the fast-continuous spreading phase, we found that all the double logarithmic plots are almost linear except for Log(ΔS)-Log(T) at higher FN concentrations. The slopes or power law exponents for frequency shift (Δf_A_) during this phase (a_2_, Δf_A_) are between 1.02-3.32 for Log(Δf_A_)-Log(T), showing no significant dependence on FN concentration; whereas for KRGD, the power law exponent (a_2_, Δf_A_) increases with the increase of RGD concentration, which is consistent with the previous reported results that the rate of spreading increases with the density of substrate ligand RGD.^[39]^ For RGD, the power law exponent for G″ (a_2_, G″) also increases with the increase of RGD concentration. The much better linear relation for Log (ΔS)-Log (T) with RGD can be ascribed to the shorter distance between cells and the RGD-modified substrate and the specific self-assembled structure of the RGD-PEG which provides higher sensitivity and stability to cells generated force ΔS (Fig. S-4). The duration of the whole second phase is between 22-84 min, showing no significant differences between FN and RGD and among different concentrations of either FN or RGD.

During the third slow spreading phase, Δf_A_ only increased slightly with time with average slope ratio over that of the second phase (a_3_/a_2_) of 6.9% and 19% for FN and RGD modified substrates, respectively. Again, the RGD modified substrates showed much better overall correlation factors (R^2^) compared to the FN modified ones; however, they are still worse than those of the second phases. The small slopes and R^2^ values are strong indications of the periodic protrusion-retraction phase.^[40]^ The cell adhesion dynamics is generally assayed by looking at the cell radii or spreading area, here we use Δf_A_ to represent the spreading area, and we found both G′ and G″ showed good adhesion dynamics, in particular G″ demonstrated easily distinguishable adhesion phases, as the addition of cells, the QCM detects the viscous and/or viscoelastic responses of the cells well ahead of their contact to the QCM surface and establishment of focal adhesion. However, the G′ characteristics of the cells can only be detected when the cells make good attachment to the QCM surface. Unlike G′ and G″, ΔS is a vector parameter, both compressive and tensile stresses may coexist, therefore, ΔS showed the worst adhesion dynamic, on the other hand, to get a complete description of the cell mechanical changes, it is important to measure all the three mechanical parameters G′, G″ and ΔS.

During the initial period of cell adhesion (0.2 h for FN, 0.5 h for RGD), we observed both increase and decrease in ΔS, suggesting co-occurrence of both contractile and protrusive forces before ΔS monotonically increased in the second phase, which is consistent with previous report.^[41]^ Furthermore, we found obvious negative ΔS that suggests protrusive force dominated, and the negative ΔS occurred on the surface coated with either FN at below or above the optimal concentration of 20∼30 µg/mL, or RGD at above the optimal concentration of 50 µg/mL. During this initial period of cell adhesion, the cell′s viscoelastic parameters changed very little, suggesting no noticeable focal adhesions or stress fibers were formed.^[39]^

In Figure 5A, the cells′ hysteresivity coefficient, η=G″/G′=tanδ was plotted as a function of cell adhesion time, which demonstrates the relative contribution of G′and G″, in addition to that both G′ and G″ increased during cell spreading. Once coming into contact with the QCM surface, the cells quickly change from a viscous liquid state (η>>100) to a viscoelastic gel state (η<10) within the first 1 h, then to a more solid-like state as the cells further spread. η is also FN concentration dependent, and it changes with time similarly as G′and G″ do. However, η remains greater than 2 in any case, which is consistent with the fact that G″ turns to increase with f more than G′ at f over 1 Hz, and becomes very close to G′ at f from 100 Hz to 1 KHz.^[42,43]^ Hence, it is reasonable that at 9 MHz in our study, the cells exhibit predominantly viscous behavior with G″ being much greater than G′.

With this technique to quantify five biomechanical parameters of the cell including elastic and viscous moduli (G′ and G″), stress exerted on the substrate by the cell (ΔS), frequency shift due to cell spreading (Δf_A_), and hysteresivity coefficient (η), we can unprecedentedly analyze the time-dependent characteristics and correlations of these parameters during cell adhesion. Fig. 4B shows representative results of G′, G″, ΔS and Δf_A_ versus time (t), and each parameter is normalized to the average value of it during the first 2 h of HUVECs adherence with 20 µg/mL FN. The results indicate that all the 4 parameters increase with time during the initial period, in particular, G′, G″ and Δf_A_ increase with time almost at the same rate. The final and steady-state ΔS, G′, G″ are in linear relationships with Δf_A_ when the substrate is coated with either FN or RGD (Figure S5). Although the relationships between G′, G″ and ΔS during adhesion of HUVECs seem to be dependent on the concentration of FN or RGD in a complicated manner (Figure S6 and Figure S7), there are common features associated with the changes of these parameters during the initial periods of cell adhesion in relation to the ligand concentration. They include: 1) ΔS increases with little increase in G′, G″ (30 µg/mL FN); 2) negative protrusive force increases with little increase in G′, G″ (75 µg/mL RGD); 3) G′ and G″ increase with little change in ΔS (0 and 40 µg/mL FN). Subsequent to the initial period of cell adhesion, the protrusive force emerges to dominate on substrate of either naked Au or FN coating created at 40 µg/mL and RGD coatings at most concentrations tested. This can be ascribed to the lack of focal adhesion formation due to either: 1) no ligands available on substrate of naked Au, or 2) the FN or RGD ligands available are too close to allow specific interactions with integrins;^[44]^ or 3) the RGD ligands available are in short chains that may be embedded in the long chains of TGME (Figure S4).

**Figure 4.**
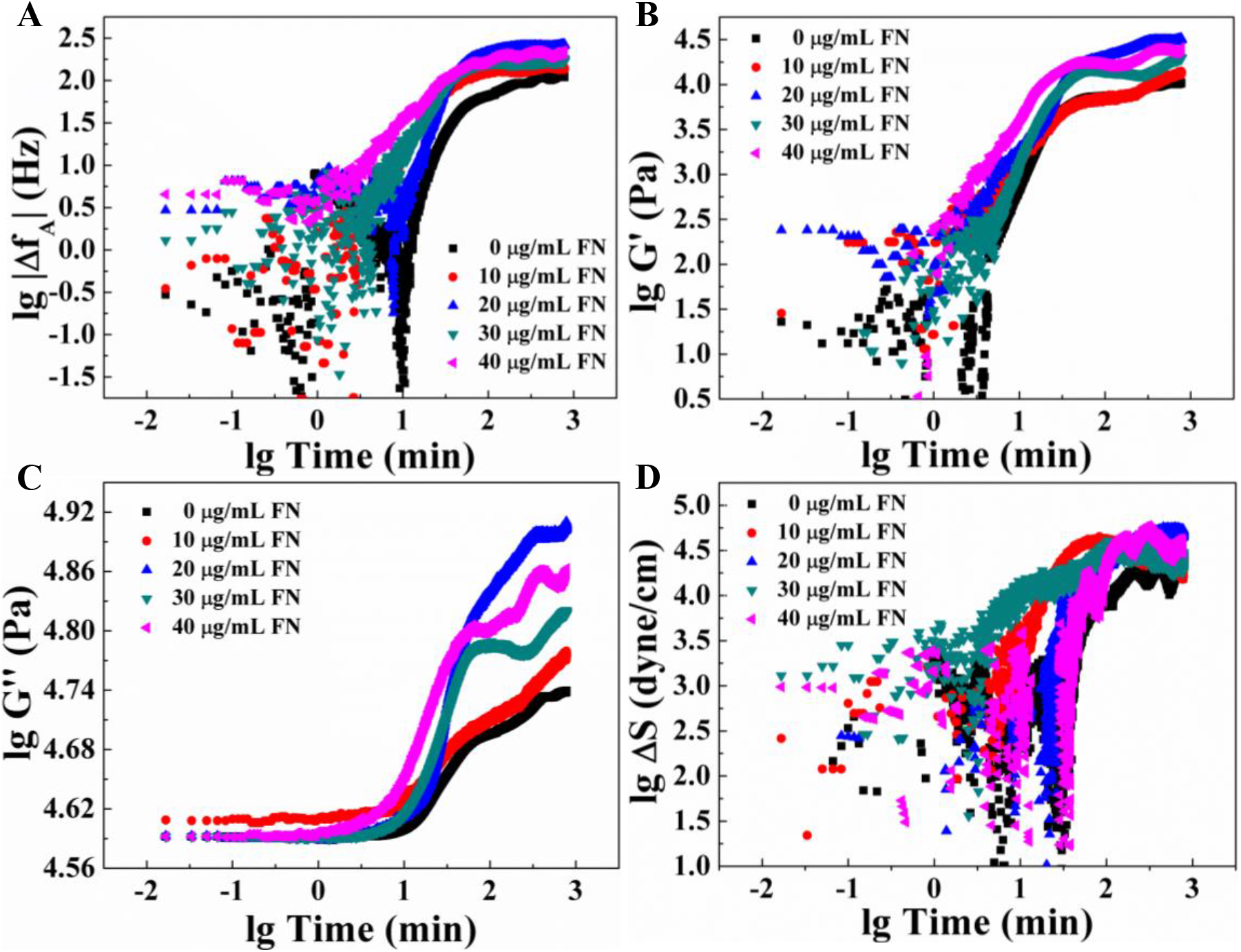
Time dependent double logarithmic plots of cytomechanical parameters during the adhesions of 20,000 HUVECs to FN modified electrodes obtained by DRPC. (A) Δf_A_, (B) G′,(C) G″, (D) ΔS.

After the initial period of cell adhesion, the cells′ mechanical parameters change with time more steadily, and ΔS becomes directly proportional to G′ or G″ obeying the cellular tensegrity model. This period may correspond to the formation of stress fibers and focal adhesions, thus resulting in greater viscoelasticity (Figure 4C). At the optimal FN concentration (20 µg/mL), the results can be fit in tensegrity model at most time points except at 0.83 h where are two distinct regions with different slopes crossing. To be comparable with other previously reported tensegrity-based experimental and theoretical results,^[45,46]^ we converted the total ΔS induced by 20000 HUVECs into a traction force by an individual HUVEC with unit of pascal by P_ECM_ (pascal) = 0.1ΔS(dyne/cm)/20,000t_cells_, where t_cells_ is the thickness of the HUVECs which was assumed to be 2 µm, and also converted the G ′ measured at 9 MHz to that measured at 0.1 Hz via oscillatory magnetic twisting cytometry,^[46]^ based on the power-law structural damping model with G′_f_ =G_0_(f/f_0_)^α^ where G_0_ and f_0_ are scale factors for modulus and frequency,^[34]^ respectively, and α is the power-law exponent usually assigned in 0.10-0.16. When α is assigned as 0.12 and 0.14, the steady G′ of HUVECs at 0.1 Hz is about 3.5 and 2.4 KPa respectively, which are within the 1-5 KPa range measured by other techniques.^[47]^

If the maximal spreading area of an individual HUVEC is 3200 µm ^2^,^[48]^ the traction force generated by the cell is estimated (F=63.84*10^−4^ΔS(dyne/cm)/20,000) to be up to 176 nN during cell adhesion on the substrate with FN coating was created at the optimal concentration. Such level of traction force is about 5 times less than that reported by others.^[48]^ We attribute this to the much stronger exponential decay of F into the metal layer before reaching the sensing interface of the quartz crystal. ^[49]^

From G′_0.1Hz_-versus-P_ECM_ relations, we calculated the slopes for the two linear regions, which turned out to be 3.58 and 1.13 when α=0.14. The lower 1.13 slope is very close to 1.2 predicted by the affine tensegrity model.^[46]^ The much larger 3.58 slope can be mainly ascribed to the underestimation of cytoskeletal prestress P due to: 1) the neglected relative contribution of microtubules P_Q_ depending on the degree of cell spreading;^[46]^ 2) the dissipation of the contractile force;^[50]^ 3) the decay of cell generated stress into the depth Z of the hard substrate.^[49]^ Moreover, the slope and linear region of the G′-versus-P_ECM_ relation also depend on the FN ligand concentration (Table S3). The G′-P_ECM_ slope (in the first region) decreased with different concentrations of FN in the order of: 20 µg/ml>30 µg/ml >40 µg/ml> 0 µg/ml >10 µg/ml. In the case of RGD coating, the highest G ′-ΔS slope occurred at 75 µg/ml, and the slope value is 1.656 that is close to half of that at 20 µg/ml FN but with a wider linear region. So far, there are only limited experimental results reported in the literature about the G′ vs. P relations with mainly linear stiffening trends and various slopes.^[45,46,51]^ Most of these studies measured the viscoelastic properties and the contractile prestress in separate experiments using different cell samples except in one case AFM and TEM were combined to measure the same single cell but not at the same time and not at the same location of cellular force sensing interface.^[52]^ Moreover, these tests are end point assays, not in a dynamic and continuous way. The method we presented here allows to measure ″passive″ viscoelastic properties and ″active″ contractile prestress simultaneously for the same population of cells from the most sensitive area of the ventral side of the cells where focal adhesions are formed. Therefore, we were able not only to capture the linear regions conforming the tensegrity model of cell mechanics and examine the effects of the ligand density on the slopes of G′ vs. P relations, but also to reveal the nonlinear G′vs. P relations. After this linear region, there is a pulse or oscillation region corresponding to the protrusion-retraction phase governed by actin polymerization and myosin contraction marked typically by a spiral rise of G′, G″ with variable net changes in the cell-induced stress.^[39]^ Figure 3D presents the relationship between G′ and G″ over the entire adhesion process of HUVECs on substrates coated with FN at different concentrations. Clearly, G′ and G″ can be characterized by nearly linear relations in different regions depending on the FN concentration. Nevertheless, the slopes of the linear relationships between G′ and G″ at different FN concentrations are not significantly different. Thus, G′ versus G″ relationships can be characterized by a unified linear relationship with a slope of 1.21 as shown in Figure 4D.

### 3.2 Effect of cell (seeding) density on cell adhesion and cytomechanical dynamics

We measured and compared the 9 MHz AT and BT cut QCM responses (Fig. S8), cells generated forces and viscoelastic moduli of 10,000, 50,000 and 100,000 HUVECs during their adhesions to 20 µg/mL FN modified gold electrodes.

Cells generate mechanical forces (traction forces) while interacting with the extracellular matrix or neighboring cells. Therefore, the total stress of a population of cells exerted is determined at least by the following factors: 1) cell number, 2) cell-matrix interactions which are determined by extracellular matrix composition and extracellular ligand density, 3) cell spreading area, 4) cell-cell interactions. For isolated single cells without cell-cell interaction, the total stress would increase with increasing cell number. As the increase of the seeding density and cells become in contact each other, the total stress exerted would decrease and even change direction from positively contractile stress to negatively extensile stress.

For a population of a total number of n isolated cells without cell-cell contact, the total stress ΔS exerted by the cells is proportional to the cell number n:

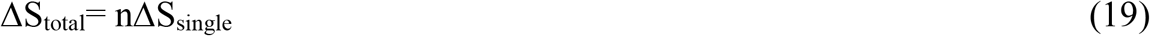

When the cells are in contact, a single cell is surrounded by its neighboring cells with total cell-cell intercellular force of ΔS_cell-cell_, then the total stress of n cells under certain cell-cell interactions at time t can be described as:

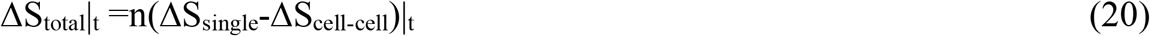

Figure 6A shows the total stresses of 10,000, 50,000 and 100,000 HUVECs exerted on the restricted Au area of ca. 0.20 cm^2^ during their adhesions to the DRPC chips, these three cell densities correspond to the three pattens of the cells: isolated, low-density confluent and high-density confluent. One can see that at the middle seeding density of 50,000 cells, the cells exerted the highest total stress after seeding the cells for 4 h. The cells’ steady-state tractions are in the following order: ΔS_50,000_> ΔS_10,000_>ΔS _100,000._ Our measured total stresses 4 h post seeding of three cell densities are consistent with previous studies which showed that the measured steady traction forces of HUVECs were in the order of low density confluent> sub-confluent>high density.^[53]^ Moreover, the DRPC technique described here allowed us to track the dynamic traction forces during the adhesions of three cell densities including the initial 4 h. Our earlier work demonstrated that QCM’s Δf and ΔR versus bovine aortic endothelial cells (BAEs) number plot 1 h post seeding exhibited a sigmoid curve shape suggesting cell-cell cooperativity in the initial cell adhesion and spreading processes;^[54]^ whereas this cell-cell cooperativity lost 24 h post seeding as a similar plot exhibited a hyperbolic curve shape. The initial 4 h of ΔS responses of Fig. 5A can be explained by the strong cell-cell interactions of high cell seeding densities. Through a contact inhibition of locomotion model,^[55]^ it was predicted that contact with other cells would decrease cells-substrate forces, and generate propulsion forces align away from neighbors.

**Figure 5.**
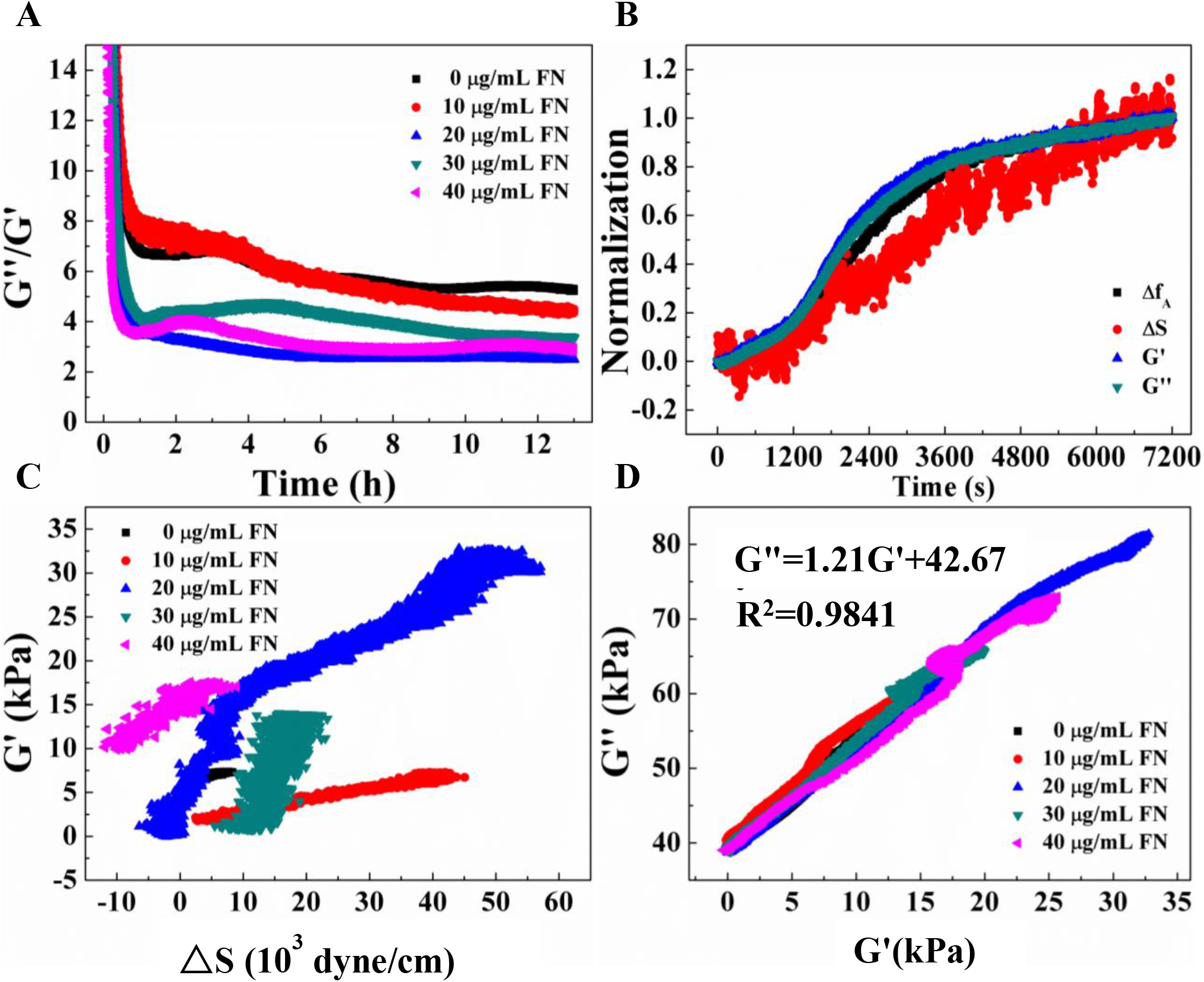
Correlations among cytomechanical parameters during the adhesions of 20,000 HUVECs on FN modified Au surfaces created from different FN concentrations.

**Figure 6.**
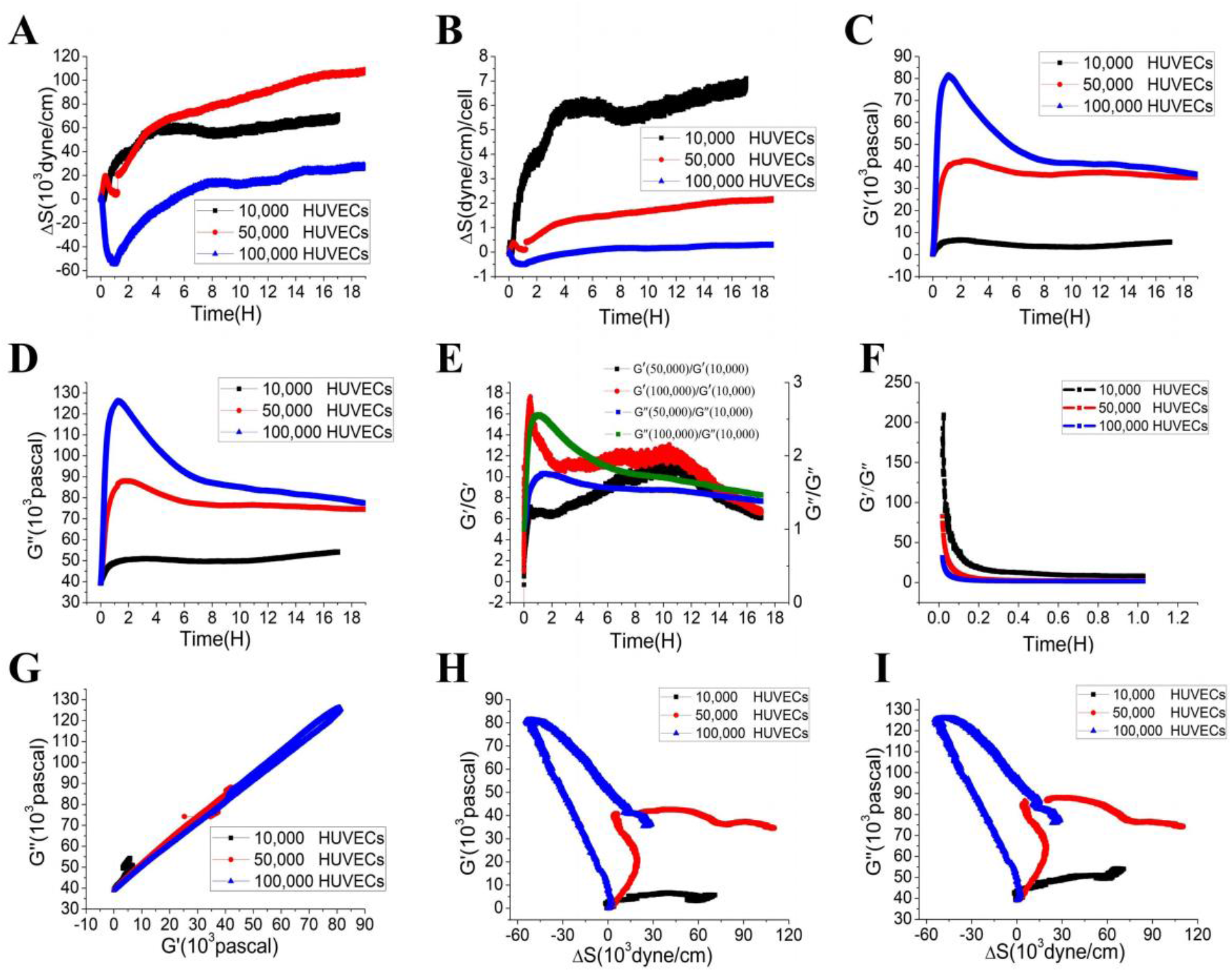
Dynamic changes of mechanical parameters of different seeding densities of HUVECs during their adhesions to 20 µg/mL FN modified quartz crystals and their correlations. (A) total exerted stress, (B) average stress exerted by a single cell, (C) storage modulus, (D) loss modulus, (E) modulus ratio, (F) loss tangent δ, (G) G′∼ΔS relations, (H) G″∼ΔS relations, (I) G″∼G′relations.

During the initial 4 h of adhesions, although 50,000 cell number is much larger than 10,000, however, because strong cell-cell interaction existing in 50,000 cells in particular during the initial 2 h, the total stress exerted was smaller than that exerted by 10,000 cells, i.e., ΔS_50,000_< ΔS_10,000_; until at 3 h, ΔS_50,000_ = ΔS_10,000_, since then ΔS_50,000_> ΔS_10,000_. The dynamic changes of ΔS at different seeding densities reflect the dynamic changes of cell-substrate and cell-cell interactions, different degrees of cell-cell interactions have antagonistic effects on cell-substrate interaction, and the stronger the degree of one kind of interaction, the weaker the degree of the other kind of interaction. For even higher density of 100,000 HUVECs, during the initial period, strong cell-cell intercellular interactions made their behaviors as an tensile monolayer or even multilayer structures and ΔS became negative;^[56]^ the tensile stress became maximum at 1 h post cell seeding, as cells spread to the QCM chips and established cell-substrate traction force weaking cell-cell interactions, the total tensile stress decreased and eventually changed to the dominated contractile stress 4 h post seeding.

Figure 6B shows the average stress exerted by a single cell of the three seeding densities. Overall, the stress exerted by single cells on the DRPC chips decreased with increasing number of the cells. In particular, the single cells within 10,000 HUVECs without cell-cell contacts had much higher ΔS_single_, their steady values reached 6-7 dyne/cm, compared to only 1-2 dyne/cm and less than 0.5 dyne/cm for 50,000 and 100,000 HUVECs respectively. Again, for 100,000 HUVECs, during the initial 4 h of cell adhesion, ΔS_single_ is negative and the cells exerted tensile stress on the chips. These results support the notion and facts that single cells are contractile and tissues are under tension that this tension increases with cell density.

Compared to vector force, cells’ viscoelasticity is a scalar parameter. The viscoelasticity of a population of cells is determined by two factors, cell-cell interaction and cell spreading area, the stronger the cell-cell interaction and the larger the average spreading area of a single cell within the cell population; the larger the cells’ viscoelastic moduli. Figures 6C and 6D showed that during the initial 1 h adhesion, the viscoelastic moduli G′and G″ increased monotonically with the increase of the seeding number of HUVECs due to increased cell-cell interactions, and with the adhesion time owing to the increased spreading area. Then for 10,000 and 50,000 HUVECs, G′ and G″ continued to increase slightly up to ca. 2.5 h followed by some drop, and the drop degree was larger for 50,000 HUVECs, and finally became stable. However, for 100,000 HUVECs, after the initial 1 h of fast increase, G′ and G″ decreased dramatically up to ca. 8 h then dropped gradually. The cells probably formed multi-layered structure initially at this high seeding density of 100,000, then the cells had to pack up with smaller sizes and reduced their spreading areas within the limited electrode area of 0.2 cm^2^ to form the monolayer structure, leading to the overall reduced viscoelasticity, eventually reached to a value close to that of the 50,000 HUVECs. It was experimentally measured that the multilayered human mammary epithelial cells (HMECs) had much higher stiffness compared to single and confluent HMECs.^[57]^ Our earlier work demonstrated via QCM that cell-cell cooperativities were maintained within 1 h of cell adhesions for different seeding densities of bovine aortic endothelial cells (BAEs).^[54]^ This may explain the initial G′ and G″ responses post 1 h seeding of HUVECs as cell-cell interactions were maintained; G′,G″ increased with increasing HUVECs number and rapidly with spreading time. Soon after 1 h seeding, G′, G″of 100,000 HUVECs dropped suddenly possibly because the cells could not maintain good contact each other owing to the too crowding cell population.

It was experimentally reported that the stiffness of a cell monolayer was 2 orders higher than the isolated cells.^[58]^ Recently, a theoretical model considering the geometric constraint effect was proposed to explain why the stiffness of collective cells are different than isolated cells and it was predicted that the elastic stiffness of the monolayer cells (E_m_) is 12.5 times of that the isolated cells (E_i_), and this ratio decreases as the increase in the thickness of the monolayer.^[59]^ The experimental results with HMECs indicated that the thickness values in the order of single, isolated cells> confluent cells > mature epithelial layers.^[57]^

Our results here (Figure 6E) showed that at 100,000 HUVECs when the cells could form a compact monolayer, the ratio of its storage modulus to that of the 10,000 isolated cells (G′_10_/G′_1_) first experienced a fast rising to over 17 owing to the possible multilayer structure, followed by the sudden drop and G′_10_/G′_1_ quickly dropped to around 12 which was maintained for over 10 h supporting the theoretical model,^[59]^ then G′_10_/G′_1_ decreased gradually and finally to 6.58 at 17.02 h possibly due to the retraction of the cells accompanied with increasing height of the monolayer.^[57]^ The lower density of 50,000 HUVECs had smaller G′_5_/G′_1_, with initial peak only about 7, then increased gradually to about 10 at 10 h approaching that of the G′_10_/G′_1_ thereafter. Compared to the effect of storage modulus, the ratios G″_m_/G″_i_ are about 1/3 of those G′_m_/G′_i_. G″_10_/G″_1_ and G″_5_/G″_1_ reached their maxima of 2.56 and 1.76 at 1.2 and 1.3 h respectively then gradually decreased to 1.46 and 1.38 at the end of 17.02 h.

Figure 6F shows the loss tangent δ=G″/G′ versus time curves during the initial 1 h adhesions of three cell densities of HUVECs and indicates that all δ decreased with time as exponential decay curves, i.e., as the cells made contact with the DRPC chips, cells became more solid structures. And as the increase of cell density, the initial δ value decreased indicating that the higher the cell density, the more solid like properties of the cells. The fitted results indicated that the higher the cell density, the higher the decay rate as evidenced by the fitted decay factors of 27.81, 30.67, 33.56 respectively. he final δ values at 17.02 h were 9.40, 2.12, 2.09 respectively, again indicates that the isolated cells are more liquid like compared to cellular monolayers.

The G″ versus G′ relations and the linear regression results were presented in Figure 6G and Table 1. Among the 3 cell densities, 100,000 had the best linear relation over the entire cell adhesion period with a slope 1.031; for 50,000 cells, the slope is 1.137 for the entire adhesion period but with poor correlation factor; while for 10,000 cells, only during the initial 2.78 h adhesion, G″∼G′ relation is linear with a slope of 1.748.

**Table 1.** Linear regression results for cellular meachanical parameters of different number of HUVECs

| Cell number | $G' \sim \Delta S$ | | | $G'' \sim \Delta S$ | | | $G'' \sim G'$ | | |
| --- | --- | --- | --- | --- | --- | --- | --- | --- | --- |
|  | Slope | R<sup>2</sup> | Time | Slope | R<sup>2</sup> | Time | Slope | R<sup>2</sup> | Time |
|  |  |  | Region (h) |  |  | Region (h) |  |  | Region (h) |
| 10000 | 0.1006 | 0.9290 | 0.13~2.41 | 0.1803 | 0.9569 | 0.13~2.41 | 1.7476 | 0.9841 | 0~2.78 |
| 50000 | 0.9462 | 0.9869 | 0~0.284 | 1.1619 | 0.9905 | 0~0.284 | 1.1372 | 0.8763 | 0~20.35 |
|  | -1.1677 | 0.9598 | 0.32~1.22 | -1.2220 | 0.9586 | 0.32~1.22 | 1.1274 | 0.9980 | 0~2.50 |
| 100000 | -1.3908 | 0.9960 | 0~1.17 | -1.4563 | 0.9950 | 0~1.17 | 1.1031 | 0.9927 | 0~20.34 |
|  | -0.6429 | 0.9754 | 1.17~20.34 | -0.7349 | 0.9839 | 1.17~20.34 |  |  |  |

Figures 6H and 6I presents the G′∼ΔS, G″∼ΔS relations for the three seeding densities of HUVECs. At low cell number of 10,000, during the very initial adhesion period (0-0.13 h), there was little change in ΔS with the increase of G′ and G″. After that, there was a limited time region (0.13-2.41 h) when G′and G″ increased roughly linearly with increasing ΔS, then the G′∼ΔS, G″∼ΔS relations severely deviated the linearities with some oscillations. At 50,000, at the beginning stage (0-0.284 h), cell-substrate traction force was dominant and the G′∼ΔS, G″∼ΔS relations were linear. However, soon after that (0.32-1.22 h), the extensile stress generated by cell-cell interactions became dominant, and the relationships between G′,G″ and ΔS became negatively linear, i.e., G′,G″ increased with decreasing ΔS. At the later stage (1.22-20.35 h), G′, G″ only decreased slightly with continuous increase of ΔS. For 100,000 HUVECs, the G′∼ΔS, G″∼ΔS relations can be clearly divided into two regions at crossing time of 1.17 h. In the first stage (0-1.17 h), cell-cell interactions induced tensile stress dominated, and there were negative linear correlations between G′, G″ and ΔS; that is, the increase of G′, G″ was accompanied with the increase of negative extensile stress. hen the cells’ generated tensile stress decreased accompanied with the decreases of G′, G″and the cells entered the second region (1.17-20.35 h) which is partially the reversable process of the first region, however, both G′, G″ and the absolute value of ΔS decreased at much slower rate comparted to the initial fast increase period, and the G′∼ΔS, G″∼ΔS slopes of the second region are only about half of those of the first region.

As discussed in the effects of ligand density, we converted the total ΔS induced by certain cell number (n) of HUVECs into an average stress generated by an individual HUVEC with unit of pascal by P_ECM_ (pascal) = 0.1ΔS(dyne/cm)/nt_cells_, where t_cells_ is the thickness of the HUVECs which was assumed to be 2 µm, and also converted the G′, G″ measured at 9 MHz to those measured at 0.1 Hz based on the power-law structural damping model with power-law exponent α assigned to 0.14. From G′_0.1Hz_∼P_ECM,_ G″_0.1Hz_∼P_ECM_ relations, we did linear regression treatments for all the linear regions and presented the results in Table 2. For 10,000 HUVECs, the slopes are 1.55 and 2.78 respectively for G′_0.1Hz_∼P_ECM,_ G″_0.1Hz_∼P_ECM_ during the adhesion of 0.13∼2.41 h, which are typical values for isolated adhered cells obeying the tensegrity model.^[40]^ However, for 50,000 cells, during the early short time (0-0.28 h), the respective stiffening slopes of 72.9 and 178.9 were even larger than reported for suspended and mitotic cells where no focal adhesion was established. ^[51]^ In our case, strong cell-cell interactions would decrease traction forces at cell−cell contacts while increasing the cell’s tensile stress. However, both cell’s tensile and contractile stress would increase the cell’s stiffness which is another reason of unusual high stiffening slopes we observed here. Subsequent to this brief stiffening period (0.32-1.22 h), even negative stiffening slopes of -89.9 and -188.4 were observed as cell’s tensile stress became dominated. For even high cell density of 100,000 HUVECs, the cells generated tensile stress once initiated the contact with the DRPC chips, both the cells’ G′, G″ were in direct proportional to the cells’ generated stress ΔS in two regions with negative slopes of -214, -224 for the first 1.17 h and -99, -113 for the second region of 1.17-20.34 h. During the first region, cell-cell interactions were the strongest resulting in overcrowding and higher density confluent pattern or even muti-monolayers, as such the cells exerted lower total forces as a concentration of forces occurring only at the cellular periphery compared to the confluent monolayer pattern of the second region.^[53]^ Whereas the cells’ viscoelastic moduli are monotonically proportional to the seeding density, i.e., the viscoelastic moduli of the first region are larger than those of the second region. Owing to the above two reasons, the slopes of G′_0.1Hz_∼P_ECM,_ G″_0.1Hz_∼P_ECM_ relations in the first region of 100,000 HUVECs were about twice of the second region.

**Table 2.**
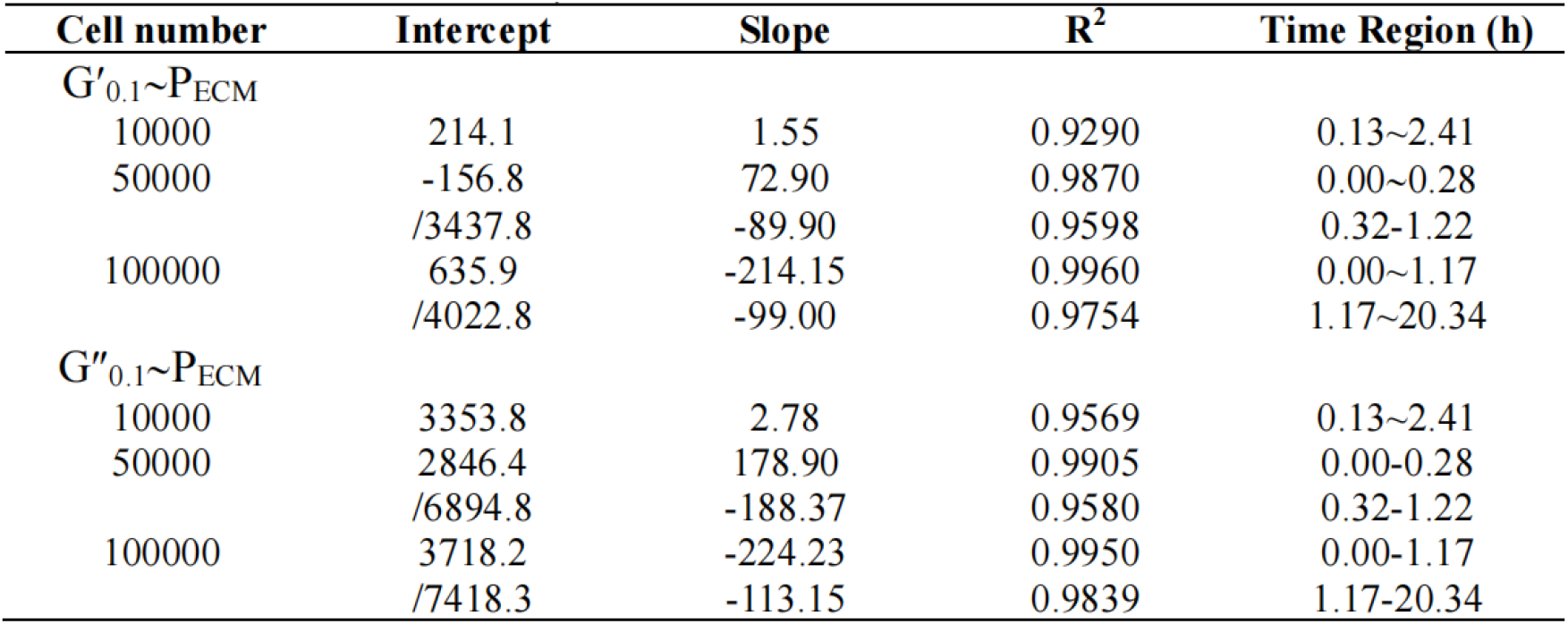
Linear regression results for G′_0.1_-versus-P_ECM_ and G″_0.1_-versus-P_ECM_ relations at different cell numbers with G′ and G″ calculated from the power-law structural damping model assuming α=0.14 and cell thickness = 2μm

| Cell number | Intercept | Slope | $R^2$ | Time Region (h) |
| --- | --- | --- | --- | --- |
| $G'_{0.1} \sim P_{ECM}$ | | | | |
| 10000 | 214.1 | 1.55 | 0.9290 | 0.13~2.41 |
| 50000 | -156.8 | 72.90 | 0.9870 | 0.00~0.28 |
|  | /3437.8 | -89.90 | 0.9598 | 0.32~1.22 |
| 100000 | 635.9 | -214.15 | 0.9960 | 0.00~1.17 |
|  | /4022.8 | -99.00 | 0.9754 | 1.17~20.34 |
| $G''_{0.1} \sim P_{ECM}$ | | | | |
| 10000 | 3353.8 | 2.78 | 0.9569 | 0.13~2.41 |
| 50000 | 2846.4 | 178.90 | 0.9905 | 0.00~0.28 |
|  | /6894.8 | -188.37 | 0.9580 | 0.32~1.22 |
| 100000 | 3718.2 | -224.23 | 0.9950 | 0.00~1.17 |
|  | /7418.3 | -113.15 | 0.9839 | 1.17~20.34 |

We failed to find negative stiffening slopes reported in the literature although tense stresses are known in monolayers and tissues whose moduli are higher than the isolated single cells.^[56,57^ Recently, nanoscale prestress and elastic modulus of a single cell were detected by AFM and negative prestress was presented for both MCF10A and MCF7 cells though no quantitative relation between elastic modulus and prestress was established because the data were too scattering.^[13]^ The DRPC technique allowed us to simultaneously measure the stresses ΔS and viscoelastic moduli G′, G″of a population of cells of different cell-cell interactions (seeding density) and obtained statistically quantitative data to reveal dynamic and complicated relations among the cellular mechanical parameters which were not captured by any other techniques.

### 3.3 Quantifying mechanical changes of HUVECs subjected to drug treatments

Figure 7 shows the effect of myosin II inhibitor blebbistatin and microtubule disruption agent nocodazole on the mechanical parameters of HUVECs measured by the DRPC technique. Upon addition of 1 µM blebbistatin, ΔS, G′, G″ all decreased significantly, which are highly consistent with the cell behaviors in response to the same drugs as detected by other conventional methods.^[60,61]^ Moreover, the increased η(G″/G′) implies that the HUVECs become a more liquid-like structure when treated with blebbistatin. Upon addition of 0.5 µM nocodazole, ΔS increased initially then turned to decrease gradually while G′, G″ increased dramatically, which are also consistent with the published results.^[62,63]^ When treated with nocodazole, however, the HUVECs become a more solid-like structure as demonstrated by the reduced η.

**Figure 7.**
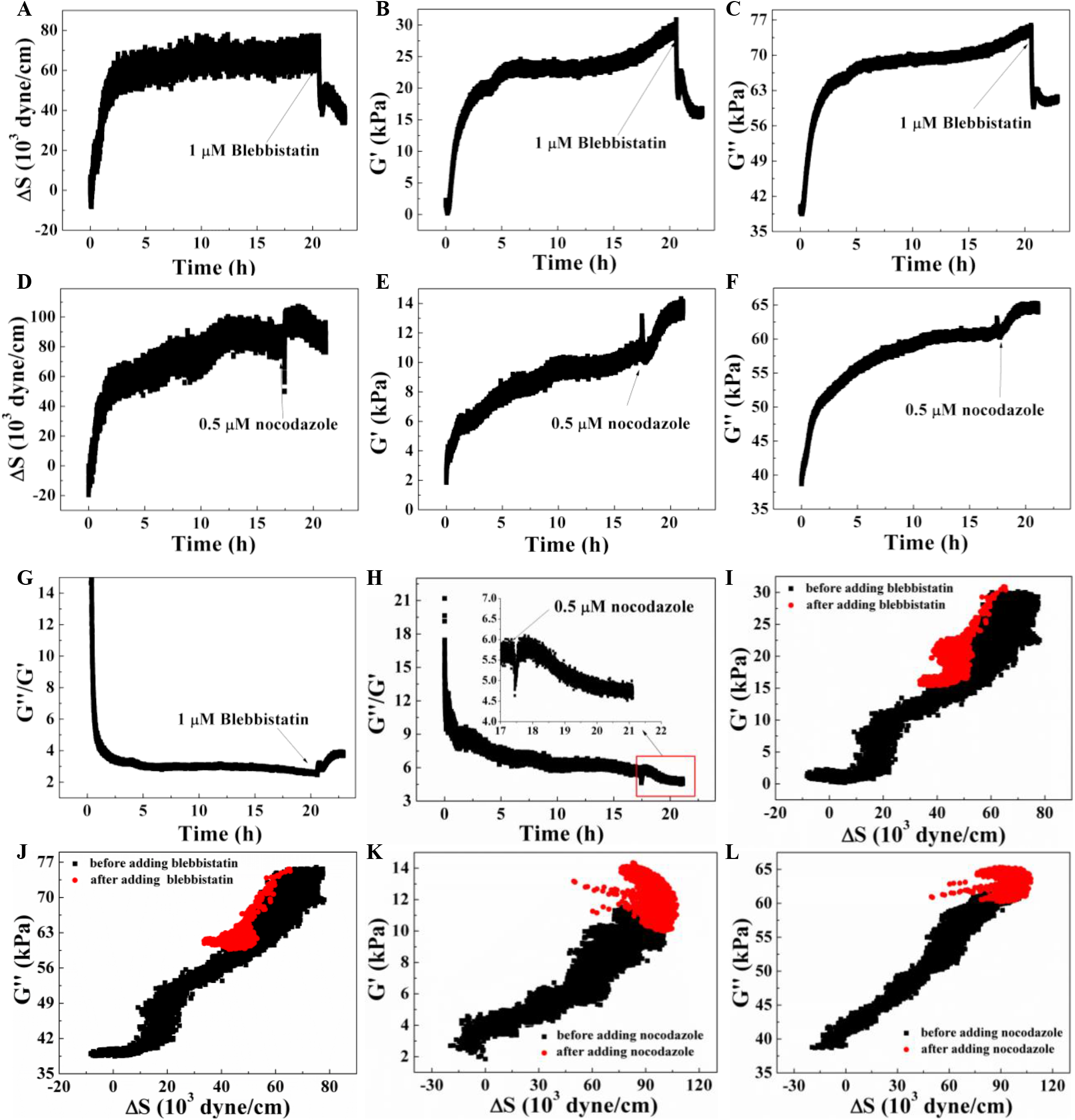
Effects of 1 µM blebbistatin and 0.5 µM nocodazole to the mechanical parameters of 20,000 HUVECs adhered to 20 µg/mL FN and 50 µg/mL RGD modified quartz crystals, respectively.

The increase in ΔS due to nocodazole may be explained by the tensegrity hypothesis that cytoskeleton-based microtubules (MTs) carry compression as they balance a portion of cell contractile stress. Upon the disruption of MTs, the portion of stress balanced by MTs would shift to the substrate, thereby causing an increase in traction. However, the treatment with either blebbistatin or nocodazole does not seem to alter the relationships between G′ and ΔS, or G″ and ΔS significantly, except for some scattered points suggesting partially destruction of the tensegrity structure which was also indicated by the reduction of ΔS when the cells were subjected to prolonged incubation of 0.5 µM nocodazole.

## 4. Materials and methods

### 4.1. Materials

Human umbilical vein endothelial cells (HUVECs) were purchased from Shanghai Bogoo Biological Technology Co. Ltd. Before the experiment, HUVECs were cultured with Dulbecco’s modified Eagle’s medium/high glucose (DMEM, Hyclone, Co., USA) containing 10% fetal bovine serum (FBS, Gibco) and 1% penicillin/streptomycin (Gibco) at 37°C and 5% CO _2_. Fibronectin was purchased from Sigma-Aldrich, and KRGD from Sangon Biotech (Shanghai) Co. Ltd. 3-mercaptopropionic acid (MPA), triethylene glycol mono-11-mercaptoundeycl ether (TGME), EDAC and N-hydroxysuccinimide (NHS) were purchased from Sigma-Aldrich. Blebbistatin and nocodazole were purchased from Sigma-Aldrich, which were dissolved in Dimethyl sulfoxide (DMSO, Solarbio). 9 MHz AT-cut and BT-cut polished quartz crystals of 0.315″square with 0.210″ diameter metal electrodes (100 Å Ti adhesion layer followed by the deposition of 1000 Å Au) were purchased from International Crystal Manufacturing Co. Ltd.

### 4.2 Modification of electrode surfaces of quartz crystals

A drop of piranha solution (30% H_2_O_2_/H_2_SO_4_, 1/3, v/v) heated at 80°C was placed on the gold electrode surface for 30 s, followed by rinsing with Milli-Q water and drying with ultra-high purity N_2_. This procedure was repeated three times. The piranha treated gold surfaces were modified with specific cell adhesion molecules RGD and fibronectin by following the procedures described in with some modifications.^[64,65]^ Specifically, the piranha treated gold surface was exposed to absolute alcohol solution containing 20 mM MPA and 1 mM TGME for 16 h at room temperature to create a self-assembled monolayer (SAM). Then the gold surface with the SAM was immersed in phosphate buffer solution (PBS, pH = 5.4) of 150 mM EDAC and 30 mM NHS for 30 min to attach the NHS group to the -COOH terminus of SAM. The substrates with NHS and PEG were sterilized with 70% ethanol for 15 min and exposed to fibronectin or KRGD at different concentrations in PBS (pH 8.2) at room temperature for 1 h. To remove loosely bound moieties from the surface after each step of surface modification, the substrate was rinsed with its original solvent and Milli-Q water, respectively.

### 4.3 Cell culture and QCM experiment

QCA 922 Quartz Crystal Analyzer with QCA 922-90 multiplexer (Seiko EG&G) was used for simultaneous measurements of 4 channels of QCM signals [66]. 400 μL DMEM media containing 20,000 HUVECs and 1% FBS was added to each well containing either FN or RGD modified QCM gold electrode substrate, and QCM frequency and resistance were continuously collected for over 13 h. For cell seeding density experiments, 9 MHz AT and BT cut quartz crystals with gold electrodes were treated with piranha solution as described above and assembled in Teflon wells, then modified with 20 μg/mL FN. 10,000, 50,000, 100,000 HUVECs were added and maintained in 5% CO_2_ incubator at 37 °C, and the QCM signals were continuously monitored during the adhesions of different seeding densities of HUVECs.

### 4.4. Effects of drugs on mechanical changes of HUVECs

Following the method described above, 9 MHz AT and BT cuts QCM gold electrodes were chemically modified with either 20 μg/mL FN or 50 μg/mL KRGD, and then assembled and enclosed into Teflon wells; 20,000 HUVECs were seeded to each well and cultured in 5% CO_2_ incubator at 37 °C for over 16 h; 10 µL media was then removed from each of the well and replaced with 10 µL media containing the chosen concentration of either blebbistatin or nocodazole to reach their final concentrations of 5 µM or 0.5 µM, respectively, and the QCM signals were continuously monitored during the drug treatments.

## Conclusions

AT-cut quartz crystals have been used as mechanical sensors for mass, lateral stress, viscosity, viscoelasticity; EerNisse established the AT and BT-cut double resonator piezoelectric technique to simultaneously measure surface mass and lateral stress of thin films. However, no theoretical work and experimental results have been reported based on QCM technique to simultaneously measure lateral stress and viscoelasticity and applied for the study of living cells. Our theoretical derivation and experimental results indicated that DRPC can quantitatively measure cells′ generated stress and viscoelasticity simultaneously for a population of cells. Here we demonstrated DRPC′s application for the study of dynamic adhesion of HUVCEs, it can also be used for the quantitative study of any other cell types of adherent cells and other cell functions such as migration, proliferation and differentiation, in particular for those functions involving collections of cells, such as wound healing as most of current quantitation methods for cell generated force are limited to single cell analysis although recently extended to cell monolayer but there is a lack of techniques to measure the total force generated by a population of cells of different seeding densities commonly used in cell culture.^[67,68]^ DRPC′s fast response time, non-invasive feature, capability of measuring multi-mechanical parameters in real-time, and compatibility with conventional culture configuration, made it ideal to serve as a common tool for cell function assays in broad biology field including cell biology and applied to disease diagnostics, therapeutics and drug efficacy assays.^[5]^ Finally, multi-well and microplates of different throughputs including high throughput DRPC chips can be made as we have demonstrated (Shen et al., 2017), which should facilitate its applications in cell phenotype assays and drug screening, that require parallel analysis of large numbers of cells.^[69]^

## Acknowledgements

This work was primarily supported by the National Natural Science Foundation of China (21275048, 31741045,11532003); and was also supported by Science and Technology Innovation Leading Plan for Hunan Provincial High-Tech Industry (2020SK2018), the Key Project of Hunan Provincial Science & Technology Department (2013TT1009), a Key Project Supported by Scientific Research Fund of Hunan Provincial Education Department (19A227). The authors would like to thank Prof. Chunyang Xiong of Peking University and Prof. Mian Long of Chinese Academy of Sciences for their constructive criticisms of the manuscript, Mr. Xiaoran Xiong of Hunan Agricultural University for his help in making figures 1&2.

## References

[1] C. A. Blair, B. L. Pruitt, Adv. Healthc. Mater. 2021, 9(8), e1901656.

[2] C. D. Pascalis, S. Etienne-Manneville, V. M. Weaver, Mol. Biol. Cell 2017, 28, 1833.

[3] B. J. Dubin-Thaler, G. Giannone, H. G. Döbereiner, M. P. Sheetz, Biophys. J. 2004, 86, 1794.

[4] P. Swiatlowska, T. Iskratsch, Biophys. Rev. 2021, 13, 611.

[5] S. Suresh, Acta Biomater. 2007, 3, 413.

[6] A. Surcel, W. P. Ng, H. West-Foyle, Q. F. Zhu, Y. X. Ren, L. B. Avery, A. K. Krenc, D. J. Meyers, R. S. Rock, R. A. Anders, C. L. F. Meyers, D. N. Robinson, Proc. Natl. Acad. Sci. USA 2015, 112, 1428.

[7] T. Iskratsch, H. Wolfenson, M.P. Sheetz, Nat. Rev. Mol. Cell Biol. 2014, 15, 825.

[8] S. Mathieu, J. B. Manneville, Curr. Opin. Cell Bio. 2019, 56, 34.

[9] D. A. Fletcher, R. D. Mullins, Nature 2010, 463, 485–492.

[10] J. Guck, E. R. Chilvers, Sci. Transl. Med. 2013, 5, 212fs41.

[11] V. Marx, Nat Methods 2019, 16, 1083.

[12] J. Guck, Biophys. Rev. 2019, 11, 667.

[13] H. X. Wang, H. Zhang, B. Da, D. B. Lu, R. Tamura, K. Goto, I. Watanabe, D. Fujita, N. Hanagata, J. Kano, T. Nakagawa, M. Noguchi, Nano Lett. 2021, 21, 1538.

[14] E.P. EerNisse, J. Appl. Phys. 1972, 43,1330.

[15] J. Y. Chen, L. S. Penn, J. Xi, Biosens Bioelectron 2017, 99, 593.

[16] L. Tan, Q. J. Xie, X. E. Jia, M. L. Guo, Y. Y. Zhang, H. Tang, S. Z. Yao, Biosens Bioelectron 2009, 24, 1603.

[17] C.T. Lim, E.H. Zhou, S.T. Quek, J. Biomech. 2006, 39, 195.

[18] R.F. Schmitt, J.W. Allen, J.F. Vetelino, J. Parks, C. Zhang, Sensors Actuat. B – Chem. 2001, 76, 95.

[19] C. Lu, in Methods and Phenomena, Vol. 7 (Eds: C. Lu, A.W. Czanderna), Elsevier, New York 1984, Ch. 2.

[20] E.P. EerNisse, J. Appl. Phys. 1973, 44, 4482.

[21] R. N. Thurston, K. Brugger, Phys. Rev. 1964, 135, 3.

[22] E.P. EerNisse, in Methods and Phenomena, Vol. 7 (Eds: Lu. v, A.W. Czanderna), Elsevier, New York 1984, Ch. 4.

[23] K. K. Kanazawa, J. G. Gordon Ii, Anal. Chim. Acta 1985, 175, 99.

[24] S. J. Martin, V. E. Granstaff, G. C. Frye, Anal. Chem. 1991, 63, 2272.

[25] H. L. Bandey, S. J. Martin, R. W. Cernosek, A. R. Hillman, Anal. Chem. 1999, 71, 2205.

[26] D. Johannsmann, Macromol. Chem. Phys. 1999, 200, 501.

[27] D. N. Clarke, A. C. Martin, Curr. Biol. 2021, 31, R667.

[28] G. N. M. Ferreira, A. C. Da-Silva, B. Tomé, Trends. Biotechnol. 2009, 27, 6897.

[29] S. I. Pyun, J. Y. Go, H. C. Shin, J. New Mat. for Electr. Sys. 2002, 5, 143.

[30] G. R. Stafford, U. Bertocci, J. Phys. Chem. C 2009, 113, 13249.

[31] J. Wang, L. M. Frostman, M. D. Ward, J. Phys. Chem. 1992, 96, 5224.

[32] L. H. Goodman, E. S. Bililign, B. W. Keller, S. G. Kenny, J. Krim, Appl. Phys. 2018, 124, 024502.

[33] a) N. Tymchenko, E. Nilebäck, M. V. Voinova, J. Gold, B. Kasemo, S. Svedhem, Biointerphases 2012, 7, 43; b) D. Johannsmann, in Bioanalytical Reviews, Vol. 2 (Eds: J. Wegener), Springer, Cham, Germany 2019, Ch. 4.

[34] H. Y. Wu, G. Y. Zhao, H. F. Zu, J. H. C. Wang, Q. M. Wang, Sensors Actuat. B – Chem. 2015, 210, 369.

[35] N. Wang, K. Naruse, D. Stamenović, J. J. Fredberg, S. M. Mijailovich, I. M. olić-Nørrelykke, T. Polte, R. Mannix, D. E. Ingber, P. Natl. Acad. of Sci. USA 2001, 98, 7765.

[36] J. Redepenning, T. K. Schlesinger, E. J. Mechalke, D. A. Puleo, R. Bizios, Anal. Chem. 1993, 65, 3378.

[37] H. G. Döbereiner, B. Dubin-Thaler, G. Giannone, H. S. Xenias, M. P. Sheetz, Phys. Rev. Lett. 2004, 93, 108105.

[38] B. J. Dubin-Thaler, G. Giannone, H. G. Döbereiner, M. P. Sheetz, Biophys. J. 2004, 86, 1794.

[39] C. A. Reinhart-King, M. Dembo, D. A. Hammer, Biophys. J. 2005, 89, 676–689.

[40] H. Wolfenson, T. Iskratsch, M. P. Sheetz, Biophys. J. 2014, 107, 2508.

[41] S. J. Henry, C. S. Chen, J. C. Crocker, D. A. Hammer, Biophys. J. 2015, 109, 699.

[42] L. Deng, X. Trepat, J. P. Butler, E. Millet, K. G. Morgan, D. A. Weitz, J. J. Fredberg, Nat. Mater. 2006, 5, 636.

[43] A. Rigato, A. Miyagi, S. Scheuring, F. Rico, Nat. Phys. 2017, 13, 771.

[44] L. Sandrin, D. Thakar, C. Goyer, P. Labbé, D. Boturyn, L. Coche-Guérente, J. Mater. Chem. B 2015, 3, 5577.

[45] N. Wang, I. M. olić-Nørrelykke, J. X. Chen, S. M. Mijailovich, J. P. Butler, J. J. Fredberg, D. Stamenović, Am. J. Physiol. Cell Physiol. 2002, 282, C606.

[46] D. Stamenović, Acta Biomater. 2005, 1, 255.

[47] M. E. Grady, R. J. Composto, D. M. Eckmann, J. Mech. Behav. Biomed. Mater. 2016, 61, 197.

[48] C. Müller, T. Pompe, Soft Matter 2016, 12, 272.

[49] S. J. He, Y. W. Su, B. H. Ji, H. J. Gao, J. Mech. Phys. Solids. 2014, 70, 116.

[50] L. Kurzawa, B. Vianay, F. Senger, T. Vignaud, L. Blanchoin, M. Théry, Mol. Biol Cell 2017, 28, 1825.

[51] E. Fischer-Friedrich, Biophys. J. 2018, 114, 419.

[52] N. Schierbaum, J. Rheinlaender, T. E. Schäffer, Soft Matter 2019, 15, 1721.

[53] A. Bajpai, J. Tong, W. Y. Qian, Y. S. Peng, W. Q. Chen, Biophys. J. 2019, 117, 1795.

[54] K. A. Marx, T. A. Zhou, M. Warren, S. J. Braunhut, Biotechnol. Progr. 2003, 19, 987.

[55] J. Zimmermann, B. A. Camley, W. J. Rappel, H. Levine, P. Natl. Acad. of Sci. USA 2016, 113, 2660.

[56] L. Balasubramaniam, A. Doostmohammadi, T. B. Saw, G. H. N. S. Narayana, R. Mueller, T. Dang, M. Thomas, S. Gupta, S. Sonam, A. S. Yap, Y. Toyama, R. M. Mège, J. M. Yeomans, B. Ladoux, Nat. Mater. 2021, 20, 1156.

[57] H. Lee, K. Bonin, M. Guthold, Biochim. Biophys. Acta Gen. Subj. BBA-Gen. Subjects 2021, 129891.

[58] A. R. Harris, L. Peter, J. Bellis, B. Baum, A. J. Kabla, G. T. P. Charras, P. Natl Acad of Sci USA 2012, 109, 16449.

[59] Y. Liu, L. Y. Zhang, B. C. Wang, G. K. Xu, X. Q. Feng, J. Mech. Phys. Solids 2020, 147, 104280.

[60] R. H. Lam, S. Weng, W. Lu, J. Fu, Integr. Biol. 2012, 4, 1289.

[61] H. Schillers, M. WaLte, K. Urbanova, H. Oberleithner, Biophys. J. 2010, 99, 3639.

[62] Y. L. Wu, W. Engl, B. Hu, P. Cai, W. R. Leow, N. S. Tan, C. T. Lim, X. Chen, ACS nano 2017, 11, 6996.

[63] Z. Al-Rekabi, K. Haase, A. E. Pelling, Exp. Cell Res. 2014, 322, 21.

[64] M. Veiseh, M. Zhang, J. Am. Chem. Soc. 2006, 128, 1197.

[65] M. Veiseh, O. Veiseh, M. C. Martin, F. Asphahani, M. Zhang, Langmuir 2007, 23, 4472.

[66] H. B. Shen, T. A. Zhou, J. J. Hu, Anal. and Bioanal. Chem. 2017, 409, 6463.

[67] W. J. Polacheck, C. S. Chen, Nat. Methods 2016, 13, 415.

[68] P. Roca-Cusachs,V. Conte, X. Trepat, Nat. Cell Biol. 2017, 19, 742–751.

[69] M. Eisenstein, Nature 2017, 544, 255.

