## Supplemental Figure 1 for "Real-Time Quantification of Cell Mechanics and Functions by Double Resonator Piezoelectric Cytometry — Theory and Study of Cellular Adhesion of HUVECs"

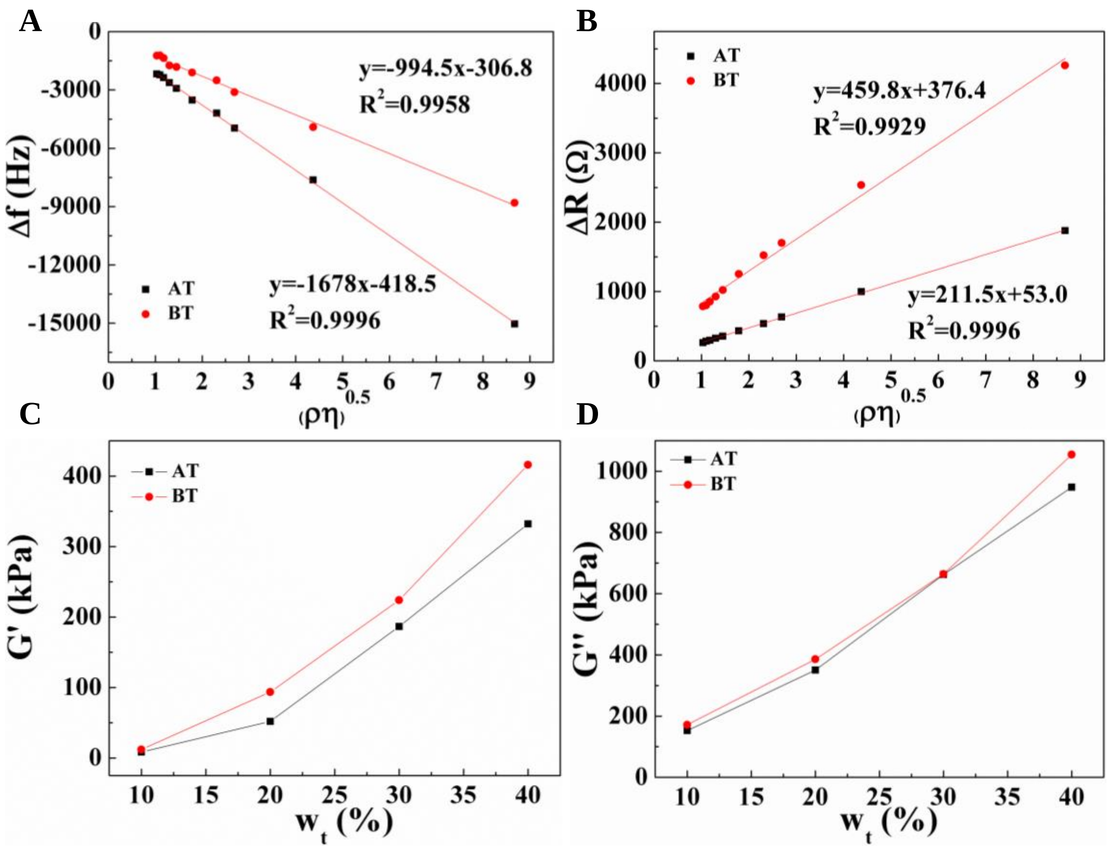


**Figure S1.** Measured results of 9 MHz AT and BT cut crystals in Newtonian sucrose and viscoelastic PEG 2000 solutions are consistent with theoretical predictions. (A, B) Measured frequency shifts (A) and motional resistance changes (B) of 9 MHz AT and BT cut crystals in different concentrations of sucrose solutions with linear fit results to the liquid $(\rho\eta)^{0.5}$. (C, D) Storage modulus (C) and loss modulus (D) of different wt% of PEG 2000 solutions at 25°C calculated from the measured frequency shifts and motional resistance changes of 9 MHz AT and BT cut crystals according to equations (15) and (16) described in the main text.
