## Supplemental Figure 2 for "Real-Time Quantification of Cell Mechanics and Functions by Double Resonator Piezoelectric Cytometry — Theory and Study of Cellular Adhesion of HUVECs"

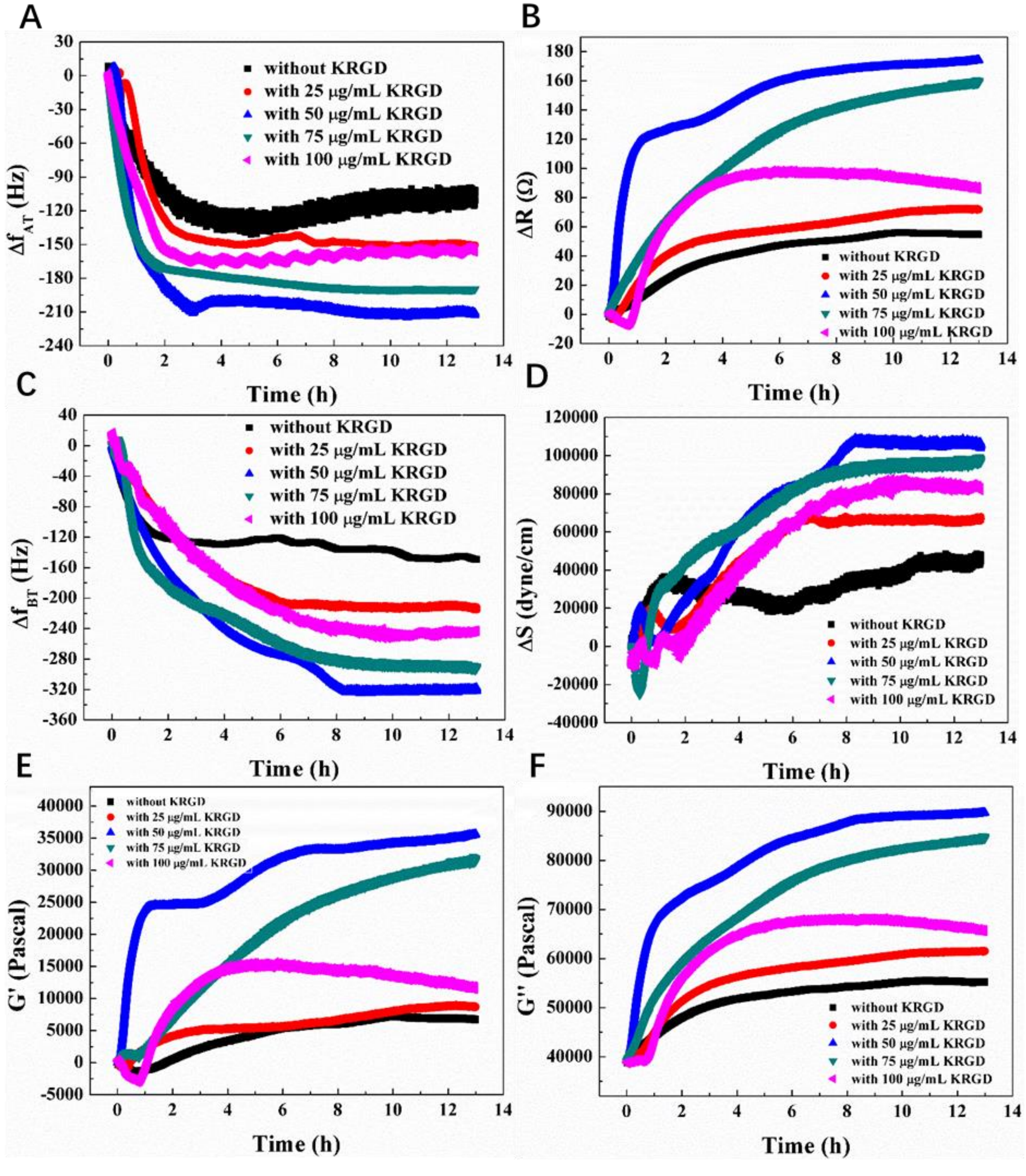


**Figure S2.** Dynamic QCM responses and cytomechanical parameters captured during the adhesions of 20,000 HUVECs on KRGD modified Au surfaces created from different KRGD concentrations. (A, B): AT cut QCM frequency shifts and motional resistance changes; (C): BT cut QCM frequency shifts; (D): Change in cells' generated stress; (E, F): Changes in cells' storage modulus and loss modulus.
