## Supplemental Figure 3 for "Real-Time Quantification of Cell Mechanics and Functions by Double Resonator Piezoelectric Cytometry — Theory and Study of Cellular Adhesion of HUVECs"

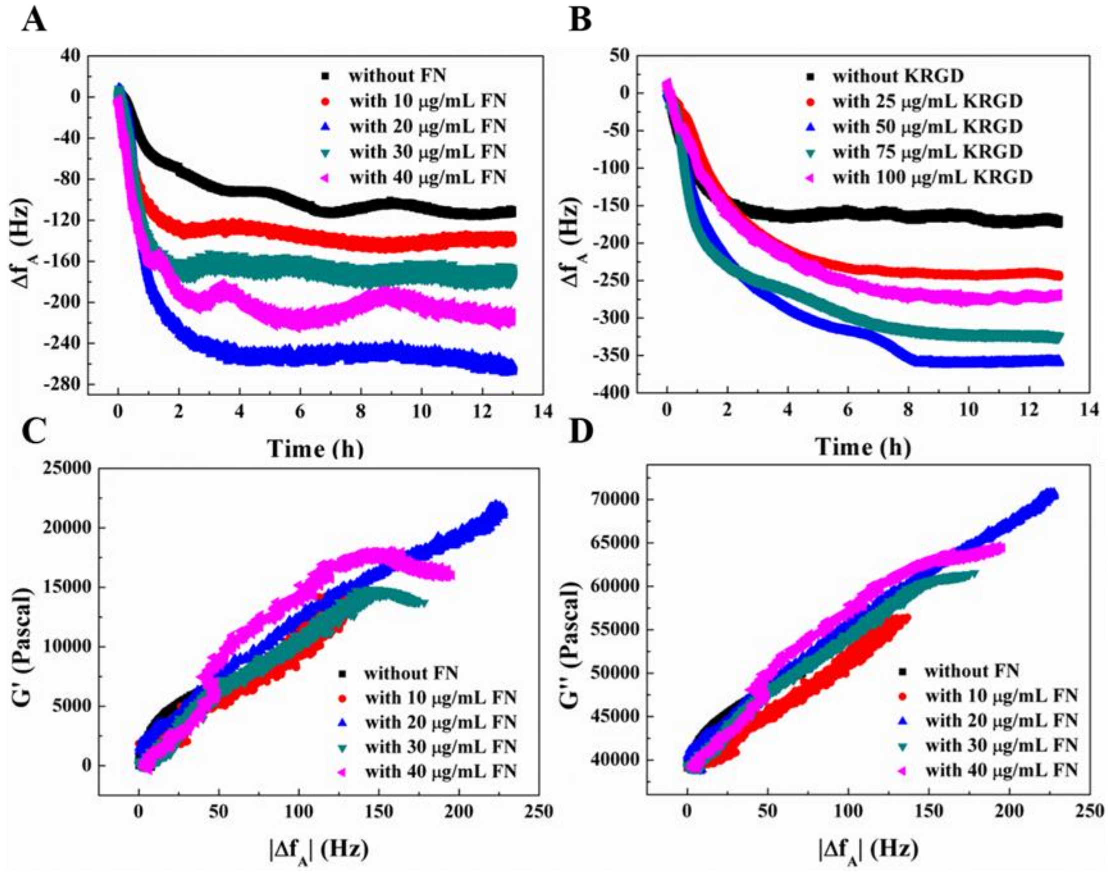


**Figure S3.** Dynamic changes in $\Delta f_A$ (frequency shifts subtracted from the surface stress induced part, indicating the degree of cells′ spreading) and G′-versus- $\Delta f_A$, G″-versus-$\Delta f_A$ relations during the adhesions of 20,000 HUVECs monitored by DRPC. (A) FN modified electrodes, (B) RGD modified electrodes. (C, D) G′-versus- $\Delta f_A$, G″-versus-$\Delta f_A$ relations during the initial 2 h of 20,000 HUVECs adhesions to FN modified electrodes. (C) G′-versus- $\Delta f_A$ relations; (D) G″-versus-$\Delta f_A$ relations.
