## Supplemental Figure 4 for "Real-Time Quantification of Cell Mechanics and Functions by Double Resonator Piezoelectric Cytometry — Theory and Study of Cellular Adhesion of HUVECs"

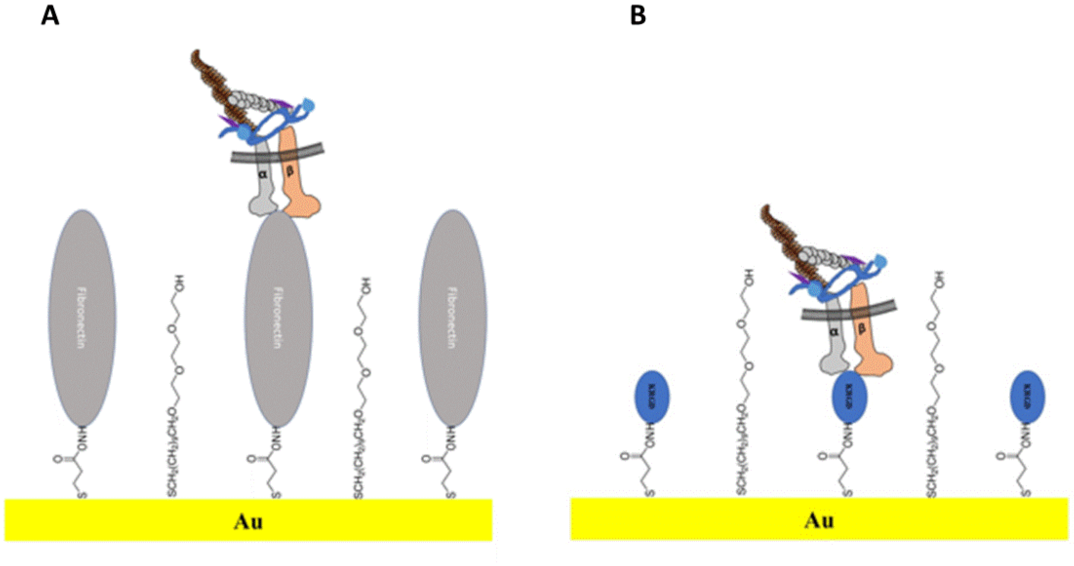


**Figure S4.** The self-assembled monolayer (SAM) created from short chain 3-mercaptopropionic acid (MPA) and longer chain triethylene glycol mono-11-mercaptoundeycl ether (TGME) may or may not affect the cell binding depending on whether protein fibronectin (schematic A) or small peptide KRGD (schematic B) is conjugated to the carboxyl group of MPA through EDAC/NHS. The transmembrane receptor integrin may have difficulty to bind ligand RGD due to steric hinderance, in particular during the initial period of the interaction, however, once bound, it may become more stable compared to integrin-fibronectin interaction as shown in schematic B.
