## Supplemental Figure 5 for "Real-Time Quantification of Cell Mechanics and Functions by Double Resonator Piezoelectric Cytometry — Theory and Study of Cellular Adhesion of HUVECs"

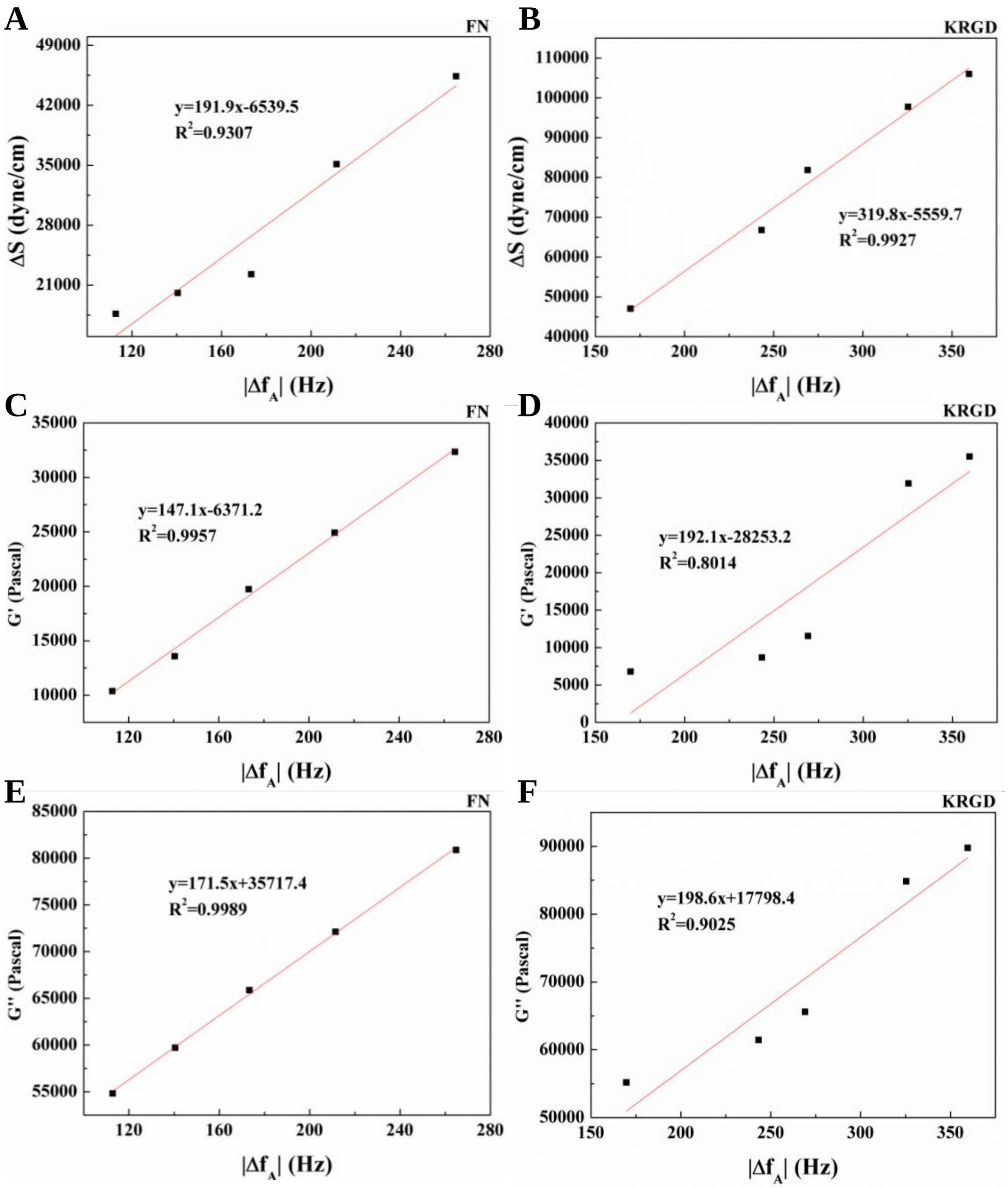


**Figure S5.** Linear relations between the final and stable cytomechanical parameters and HUVECs adhesions induced frequency shifts ($\Delta f_A$) for FN (0-40 µg/mL) and KRGD (0-100 µg/mL) modified electrodes measured by DRPC. (A,C,E): FN modified electrodes, (A) $\Delta S$-$\Delta f_A$, (C) $G'$-$\Delta f_A$, (E) $G''$-$f_A$; (B,D,F): KRGD modified electrodes, (B) $\Delta S$-$f_A$, (D) $G'$-$f_A$, (F) $G''$-$f_A$.
