## Supplemental Figure 6 for "Real-Time Quantification of Cell Mechanics and Functions by Double Resonator Piezoelectric Cytometry — Theory and Study of Cellular Adhesion of HUVECs"

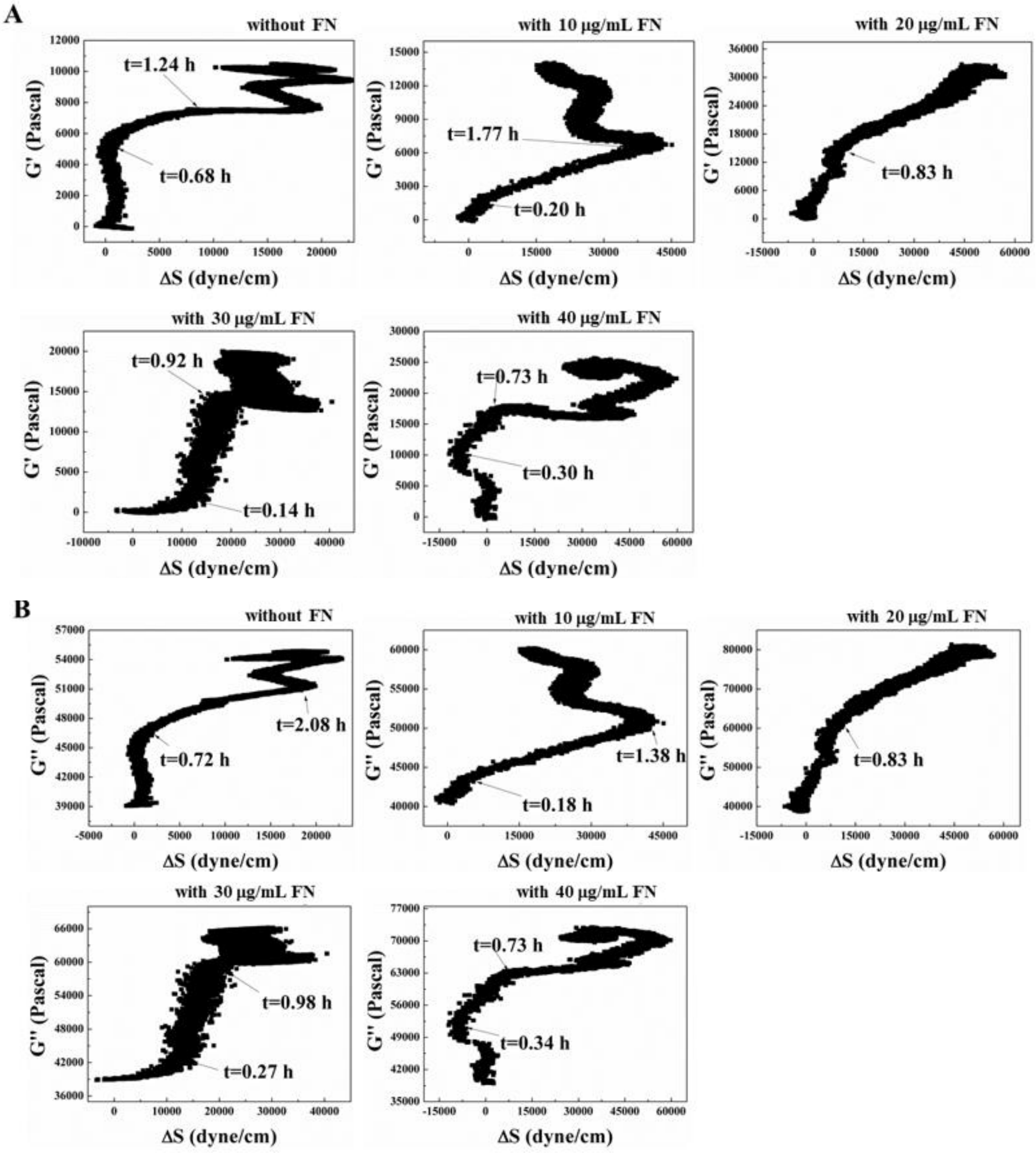


**Figure S6.** Dynamic relations between viscoelastic moduli and cells′ generated stresses during the adhesions of 20,000 HUVECs to FN modified electrodes. (A): G′-versus-ΔS relations, (B): G″-versus-ΔS relations. The time at each transition was marked by arrowhead.
