## Supplemental Figure 8 for "Real-Time Quantification of Cell Mechanics and Functions by Double Resonator Piezoelectric Cytometry — Theory and Study of Cellular Adhesion of HUVECs"

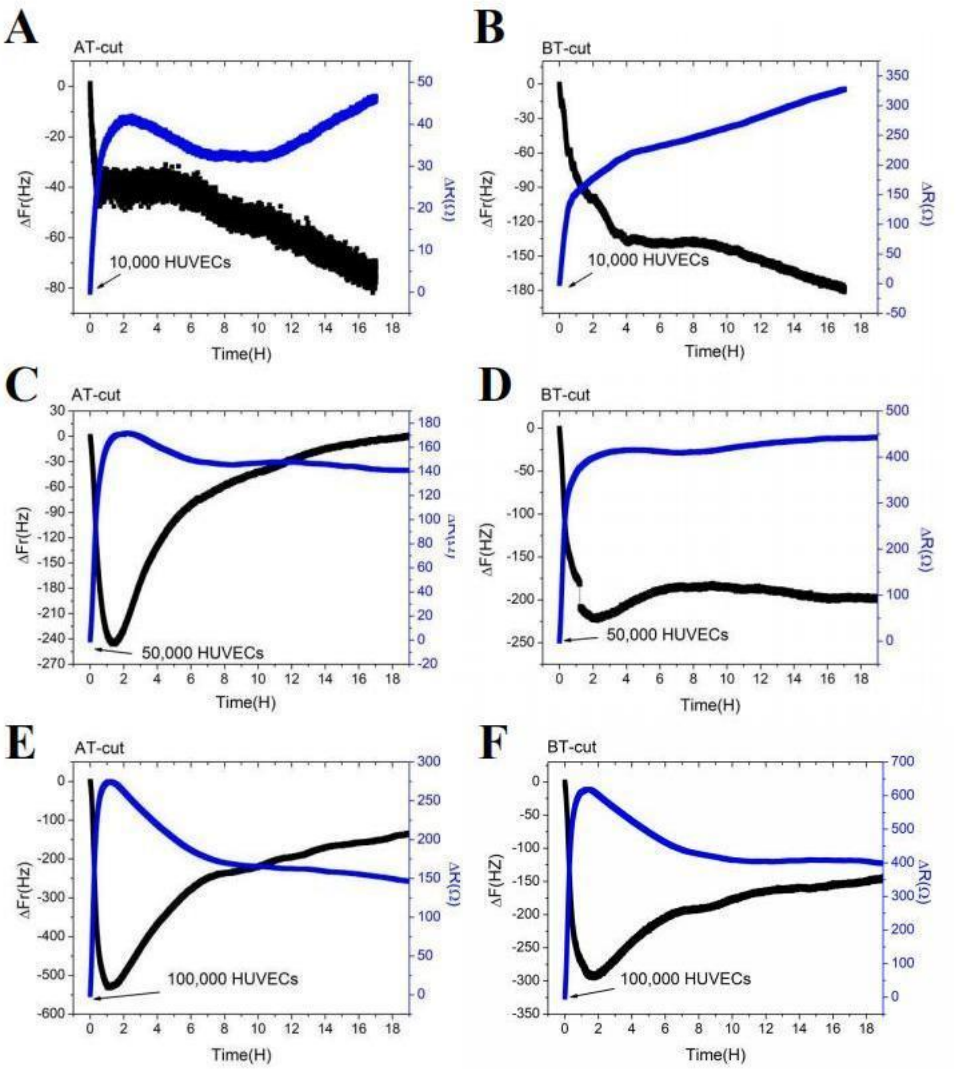


**Figure S8.** QCM responses during the adhesions of different seeding densities of HUVECs to 20 μg/mL FN modified AT and BT cut quartz crystals.
