## Supplemental Table 1 for "Real-Time Quantification of Cell Mechanics and Functions by Double Resonator Piezoelectric Cytometry — Theory and Study of Cellular Adhesion of HUVECs"

**Table S1.** Linear regression results for G′-versus-$\Delta f_A$ and G″-versus-$\Delta f_A$ relations (G′,G″:Pa; $\Delta f_A$:Hz)

| **Concentration** | **Intercept** | | **Slope** | | $R^2$ | |
|---|---|---|---|---|---|---|
| | G′-$\Delta f_A$ | G″-$\Delta f_A$ | G′-$\Delta f_A$ | G″-$\Delta f_A$ | G′-$\Delta f_A$ | G″-$\Delta f_A$ |
| FN (μg/mL) | | | | | | |
| 0 | 1685.9 | 40673.8 | 94.3 | 142.1 | 0.8925 | 0.9534 |
| 10 | 844.3 | 41552.8 | 50.5 | 79.3 | 0.9423 | 0.9913 |
| 20 | 2125.8 | 41012.6 | 87.4 | 132.6 | 0.9842 | 0.9930 |
| 30 | 1179.4 | 40220.7 | 84.3 | 131.1 | 0.9572 | 0.9859 |
| 40 | 3616.9 | 42341.2 | 84.1 | 129.4 | 0.8202 | 0.9409 |
| KRGD (μg/mL) | | | | | | |
| 0 | -5079.5 | 38070.6 | 52.0 | 59.9 | 0.8751 | 0.9480 |
| 25 | 1628.7 | 38498.1 | 35.5 | 89.3 | 0.9959 | 0.9982 |
| 50 | 10777.3 | 44852.1 | 126.9 | 139.5 | 0.9604 | 0.9373 |
| 75 | -14598.8 | 38814.5 | 103.8 | 77.5 | 0.9304 | 0.9643 |
| 100 | -14060.3 | 25562.4 | 163.8 | 194.2 | 0.9716 | 0.9872 |
