## Supplemental Table 2 for "Real-Time Quantification of Cell Mechanics and Functions by Double Resonator Piezoelectric Cytometry — Theory and Study of Cellular Adhesion of HUVECs"

**Table S2.** Linear fit results of double logarithmic plots during the fast and continuous spreading phase obtained with FN or KRGD modified electrodes ($\Delta f_A$:Hz; $\Delta S$:dyne/cm; $G'$,$G''$:Pa)

| Concentration | $lg\|\Delta f_A\|$-lgT | | $lg\Delta S$-lgT | | $lgG'$-lgT | | $lgG''$-lgT | |
|---|---|---|---|---|---|---|---|---|
| | Slope | $R^2$ | Slope | $R^2$ | Slope | $R^2$ | Slope | $R^2$ |
| FN (μg/mL) | | | | | | | | |
| 0 | 3.32 | 0.9527 | 2.19 | 0.9001 | 2.12 | 0.9969 | 0.13 | 0.9992 |
| 10 | 1.50 | 0.9797 | 1.91 | 0.9392 | 1.05 | 0.9599 | 0.10 | 0.9884 |
| 20 | 2.69 | 0.9585 | 1.55 | 0.8563 | 1.30 | 0.9657 | 0.41 | 0.9950 |
| 30 | 1.46 | 0.9572 | 0.67 | 0.6280 | 1.96 | 0.9886 | 0.25 | 0.9959 |
| 40 | 1.02 | 0.9873 | 2.98 | 0.3185 | 1.34 | 0.9906 | 0.24 | 0.9978 |
| KRGD (μg/mL) | | | | | | | | |
| 0 | 0.48 | 0.9851 | 0.84 | 0.9673 | 0.94 | 0.9517 | 0.11 | 0.9946 |
| 25 | 1.01 | 0.9702 | 1.55 | 0.9722 | 0.39 | 0.9287 | 0.12 | 0.9601 |
| 50 | 1.52 | 0.9965 | 1.26 | 0.9661 | 1.69 | 0.9895 | 0.16 | 0.9588 |
| 75 | 1.71 | 0.9927 | 0.57 | 0.9931 | 1.21 | 0.9792 | 0.21 | 0.9933 |
| 100 | 2.60 | 0.8989 | 1.64 | 0.9666 | 1.75 | 0.9610 | 0.26 | 0.9663 |
