## Supplemental Table 3 for "Real-Time Quantification of Cell Mechanics and Functions by Double Resonator Piezoelectric Cytometry — Theory and Study of Cellular Adhesion of HUVECs"

**Table S3.** Linear regression results for $G'_{0.1}$-versus-$P_{ECM}$ relations with G′ calculated from the power-law structural damping model assuming cell thickness = 2 μm ($P_{ECM}$, G′: Pa)

| Concentration | Intercept | Slope | $R^2$ | Time Range |
|---|---|---|---|---|
| FN (μg/mL) | | α=0.12 | | |
| 0 | 654.3 | 0.87 | 0.9240 | t=0.68~1.24 h |
| 10 | 177.3 | 0.56 | 0.9734 | t=0.20~1.77 h |
| 20 | 499.0 /1362.7 | 5.11 /1.62 | 0.8780 /0.9283 | t=0.00 ~0.83 h /t=0.83~13.0 h |
| 30 | -883.4 | 4.71 | 0.6104 | t=0.14~ 0.92 h |
| 40 | 1681.9 | 1.86 | 0.8253 | t=0.30~ 0.73 h |
| FN (μg/mL) | | α=0.14 | | |
| 0 | 458.0 | 0.61 | 0.9240 | t=0.68~1.24 h |
| 10 | 124.1 | 0.39 | 0.9734 | t=0.20 ~1.77 h |
| 20 | 349.3 /953.9 | 3.58 /1.13 | 0.8780 /0.9283 | t=0.00~ 0.83 h /t=0.83~13.0 h |
| 30 | -618.4 | 3.30 | 0.6104 | t=0.14 ~0.92 h |
| 40 | 1177.4 | 1.30 | 0.8253 | t=0.30 ~0.73 h |
| KRGD (μg/mL) | | α=0.12 | | |
| 0 | 693.5 | 0.36 | 0.8882 | t=5.61~1.24 h |
| 25 | 784.1 | 0.11 | 0.9228 | t=3.36~6.35 h |
| 50 | 1700.5 | 1.17 | 0.9804 | t=3.55~6.90 h |
| 75 | -891.2 | 2.37 | 0.9925 | t=1.16~7.57 h |
| 100 | 1327.0 | 0.80 | 0.9205 | t=2.44~3.84 h |
| KRGD (μg/mL) | | α=0.14 | | |
| 0 | 485.4 | 0.25 | 0.8882 | t=5.61~1.24 h |
| 25 | 548.9 | 0.08 | 0.9228 | t=3.36~6.35 h |
| 50 | 1190.4 | 0.82 | 0.9804 | t=3.55~6.90 h |
| 75 | -623.9 | 1.66 | 0.9925 | t=1.16~7.57 h |
| 100 | 928.9 | 0.57 | 0.9205 | t=2.44~3.84 h |
